# Tomato geranylgeranyl diphosphate synthase isoform 1 specifically interacts with phytoene synthase isoform 3 to produce strigolactones in tomato roots

**DOI:** 10.1101/2022.11.01.514744

**Authors:** Miguel Ezquerro, Changsheng Li, M. Victoria Barja, Esteban Burbano-Erazo, Julia Pérez-Pérez, Yanting Wang, Lemeng Dong, Purificación Lisón, M. Pilar López-Gresa, Harro J. Bouwmeester, Manuel Rodríguez-Concepción

## Abstract

- Carotenoids are photoprotectant pigments and precursors of the hormones abscisic acid (ABA) and strigolactones (SL). Carotenoids are produced in plastids from geranylgeranyl diphosphate (GGPP), which is diverted to the carotenoid pathway by phytoene synthase (PSY). In tomato (*Solanum lycopersicum*), 3 genes encode plastid-targeted GGPP synthases (*SlG1 to 3*) and 3 genes encode PSY isoforms (*PSY1 to 3*).
- Here we investigated the function of SlG1 by generating loss-of-function lines and combining their metabolic and physiological phenotypes with gene co-expression and co-immunoprecipitation analyses.
- Leaves and fruits of *slg1* lines showed a wild-type phenotype in terms of isoprenoid accumulation, photosynthesis and development. Consistently, *SlG1* is co-expressed with *PSY3* and other genes involved in the production of carotenoids and SL (but not ABA) only in roots. SlG1 was also found to physically interact with the root-specific PSY3 isoform (and not with PSY1 and PSY2). Root SL (but not ABA) levels were reduced in *slg1* lines.
- Our results confirm a specific role of SlG1 in SL production in combination with PSY3. This role appears to be restricted to roots as *slg1* plants do not exhibit the shoot phenotype displayed by other SL-deficient mutants.

## Introduction

Isoprenoids are one of the most diverse family of compounds in all living organisms, with plants displaying the highest functional and structural variation (Bouvier et al., 2005). The universal building blocks for the biosynthesis of all isoprenoids are isopentenyl diphosphate (IPP) and its allylic isomer dimethylallyl diphosphate (DMAPP). Both five-carbon (C5) isoprenoid-building molecules are produced in plants by the mevalonic acid (MVA) pathway in the cytosol and the methylerythritol 4-phosphate (MEP) pathway in the plastids (Pulido et al., 2012; Tholl, 2015). Condensation of one or more molecules of IPP to one molecule of DMAPP produces C10 geranyl diphosphate (GPP), C15 farnesyl diphosphate (FPP) and C20 geranylgeranyl diphosphate (GGPP), that are the precursors for most downstream isoprenoid compounds in different cell compartments (Ruiz-Sola & Rodríguez-Concepción, 2012; Zhou & Pichersky, 2020). In the cytosol, FPP is used to synthesize C15 sesquiterpenes and C30 triterpenes required for defensive responses, membrane structure and prenylation of proteins (Thulasiram & Poulter, 2006). In the plastids, GPP is used to make C10 monoterpenes (mostly volatile compounds related to aroma and plant-pathogen interactions) (Chen et al., 2015; Degenhardt et al., 2009) and GGPP is the precursor of gibberellins (GAs) and several photosynthesis-related isoprenoids such as carotenoids, tocopherols, chlorophylls, plastoquinone and phylloquinones (Barja & Rodriguez-Concepcion, 2021). GGPP is also used to produce C20 diterpenes in the cytosol, and both FPP and GGPP are produced in mitochondria from imported MVA-derived IPP and DMAPP for ubiquinone and C20 diterpenoid biosynthesis (Barja & Rodriguez-Concepcion, 2021; Thulasiram & Poulter, 2006).

C40 carotenoids are GGPP-derived plastidial isoprenoids that function as precursors of vitamin A and health-promoting phytonutrients in the human diet and have a great industrial interest as natural pigments (Rodriguez-Concepcion et al., 2018; Ruiz-Sola & Rodríguez-Concepción, 2012). In plants, carotenoids act as photoprotectors in leaves, as pigments in the flowers and fruits of many plant species and as precursors of apocarotenoids, including bioactive compounds such as the hormones abscisic acid (ABA) and strigolactones (SLs) (Al-Babili & Bouwmeester, 2015; Rodriguez-Concepcion et al., 2018). Despite their biological and economic relevance, the factors that integrate and coordinate carotenoid biosynthesis with plant metabolism and development remain little known. In this context understanding how GGPP is channeled to the production of carotenoids for particular functions in diverse tissues, developmental stages and environmental conditions remains a pivotal question.

In plastids, GGPP is produced from MEP-derived IPP and DMAPP by GGPP synthase (GGPPS) enzymes (Figure 1). The first committed and main flux-controlling step of the carotenoid biosynthesis pathway is the condensation of two molecules of GGPP into phytoene catalyzed by phytoene synthase (PSY). Next, phytoene is desaturated and isomerized to lycopene, and the ends of the linear lycopene chain are cyclized to form β-carotene (with two β rings) or α-carotene (with one β and one ε ring). Oxidation of the rings gives rise to xanthophylls such as violaxanthin and neoxanthin from β-carotene or lutein from α-carotene (Figure 1a) (Rodriguez-Concepcion et al., 2018).

**Figure 1.**
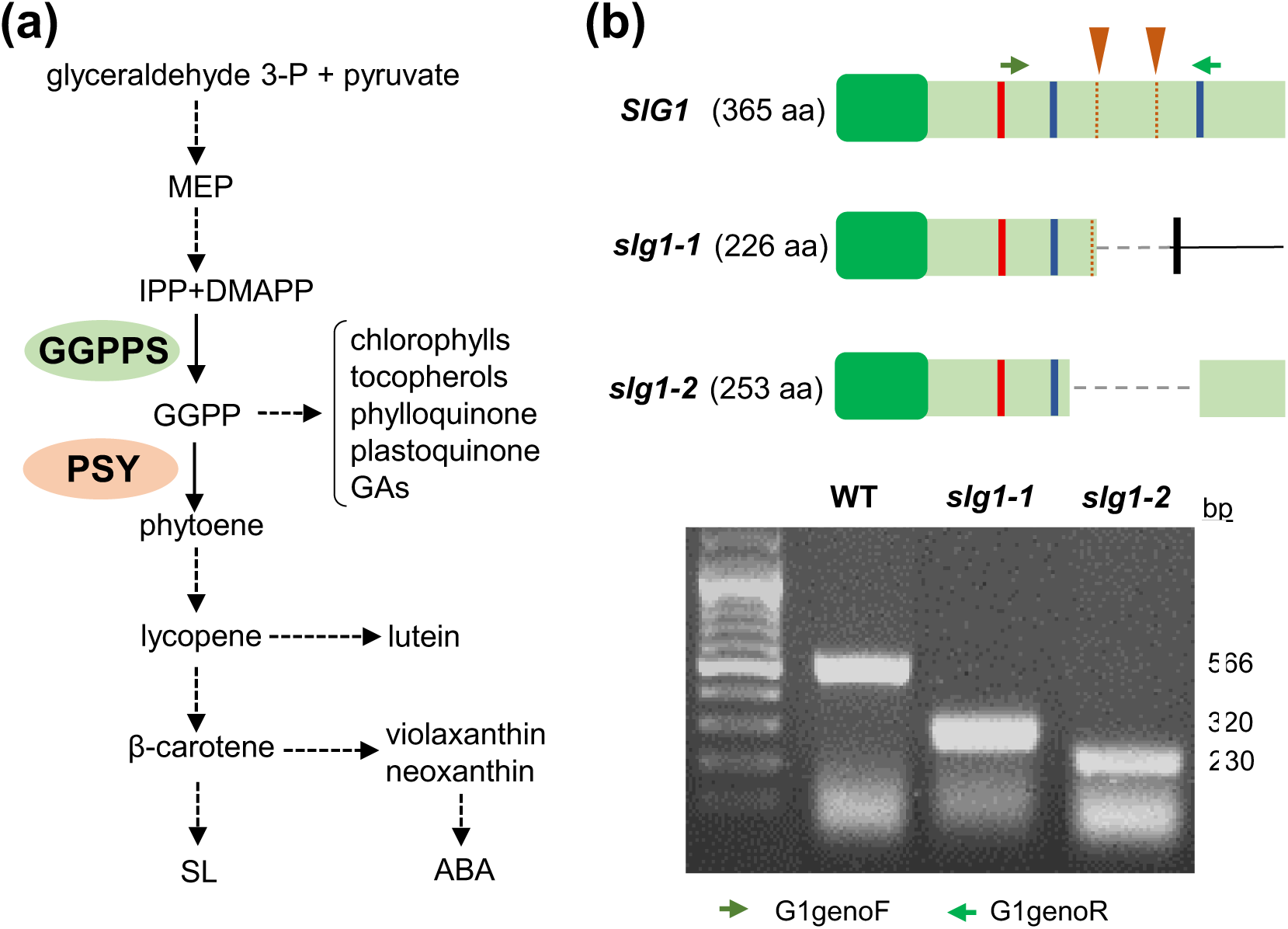
Carotenoid pathway and tomato mutants. (**a**) Carotenoid biosynthesis pathway. Dashed arrows represent multiple steps. The reactions catalyzed by geranylgeranyl diphosphate synthase (GGPPS) and phytoene synthase (PSY) are marked. **(b)** Scheme representing the wild-type SlG1 protein and the mutant versions generated in the corresponding CRISPR-Cas9-generated alleles (see Figure S1–S2 for further details). Dark green boxes represent plastid transit peptides. The regions targeted by the designed sgRNAs are indicated with orange arrowheads and dotted lines. Red and blue bars mark the position of conserved domains required for GGPPS activity (protein-protein interaction domains and Asp-rich domains, respectively). Deletions are shown with a dashed line. Green arrows represent the position of primers for PCR-based genotyping. The agarose gel shows the PCR genotyping products using these primers.

In the model plant *Arabidopsis thaliana* PSY is encoded by one gene and the resulting protein directly interacts with AtG11, the main plastidial GGPPS. This interaction likely facilitates the channeling of GGPP to the production of carotenoids (Camagna et al., 2019; Ruiz-Sola et al., 2016a; Ruiz-Sola et al., 2016b). A more complex scenario is found in tomato (*Solanum lycopersicum*). The extra roles associated with carotenoids in this species compared to Arabidopsis (including flower and fruit pigmentation and root mycorrhization) together with genome duplication events have resulted in small PSY and GGPPS gene families. With regard to PSY, tomato has an isoform with a primary role in pigmentation of flowers and ripe fruit (PSY1, Solyc03g031860), another one mainly involved in the production of carotenoids for photosynthesis and photoprotection in leaves (PSY2, Solyc02g081330) and a third one associated to SL and apocarotenoid biosynthesis in roots (PSY3, Solyc01g005940) (Ezquerro et al., 2022; Stauder et al., 2018). From the three plastid-targeted GGPPS isoforms identified in tomato, herein referred to as SlG1 (Solyc11g011240), SlG2 (Solyc04g079960) and SlG3 (Solyc02g085700) (Barja et al., 2021; Zhou & Pichersky, 2020), only SlG2 and SlG3 have been studied in detail. They both produce GGPP for carotenoid synthesis in leaves and fruits, with SlG3 being the main housekeeping isoform and SlG2 acting as a helper enzyme to meet peak demands of GGPP in both organs. SlG2 can be co-immunoprecipitated with both PSY1 and PSY2, but SlG3 cannot (Barja et al. 2021). Although SlG1 is also an active plastid-targeted GGPPS enzyme (Barja et al., 2021; Zhou & Pichersky, 2020), it cannot complement the loss of SlG2 and SlG3 in double mutants, which show an embryo-lethal phenotype like that reported for AtG11-defective Arabidopsis mutants (Ruiz-Sola et al., 2016a; Ruiz-Sola et al., 2016b; Barja et al. 2021). Indeed, *SlG1* transcripts are much less abundant than those of *SlG2* and *SlG3* in most plant tissues (Barja et al., 2021; Zhou & Pichersky, 2020). In leaves, *SlG1* expression is normally low but it is induced following spider mite feeding, wounding and elicitor treatments correlating with the production of defense-related diterpenoid volatiles (Ament et al., 2006). Other gene expression data suggest that SlG1 might have a role in roots during mycorrhization (Stauder et al., 2018; Barja et al. 2021).

Under nitrogen and/or phosphate starvation, the roots of many plant species (including tomato) exude small quantities of carotenoid-derived SL to promote recognition and colonization by arbuscular mycorrhizal (AM) fungi (Matthys et al., 2016; Stauder et al., 2018; Yoneyama et al., 2008; Zhang et al., 2014). These symbiotic AM fungi help the plant by providing water and mineral nutrients in poor soils, in exchange for carbon products biosynthesized by the plant (Bouwmeester et al., 2007; Yoneyama et al., 2008). Carotenoid metabolism is in turn stimulated in AM roots, resulting in the production of high amounts of pigments such as mycorrhadicins and other apocarotenoids including blumenols, zaxinone and anchorene that modulate the establishment of the AM symbiosis and rhizospheric interactions (Baslam et al., 2013; Fester et al., 2002; Moreno et al., 2021; Stauder et al., 2018). A coordinated role for *SlG1* and *PSY3* has been proposed for SL and AM-associated apocarotenoid biosynthesis in roots, mainly based on expression data (Barja et al., 2021; Stauder et al., 2018). *SlG1* and *PSY3* are indeed the most highly up-regulated genes encoding GGPPS and PSY isoforms when roots are mycorrhized. However, *SlG2* and *PSY1* also show increased transcript levels in mycorrhized roots compared to non-mycorrhized controls (Barja et al., 2021; Stauder et al., 2018). This, together with the observation that the basal expression levels of *SlG1* and *PSY3* in roots is lower than that of *SlG2* and *PSY1* (Barja et al., 2021; Fantini et al. 2013), suggest that more than one isoform of these two enzymes might be providing precursors for carotenoids and derived compounds in roots. To experimentally test this hypothesis and better understand the biological role of SlG1 in tomato, we created CRISPR-Cas9-edited lines defective in SlG1. Here we report their generation and characterization and demonstrate the existence of a highly specific SlG1-PSY3 interaction to produce SL in tomato roots.

## Materials and methods

**Plant material.** Tomato (*Solanum lycopersicum* var. MicroTom) plants were used for experiments. The *ccd7* line, previously referred to as *CCD7-AS*, was made by introgressing a construct expressing a *CCD7* antisense from M82 (Vogel et al., 2010) into MicroTom by successive backcrosses (Pino et al., 2022). Seeds were surface-sterilized by a 30 min water wash followed by a 15 min incubation in 10 ml of 40% bleach with 10 µl of Tween-20. After three consecutive 10 min washes with sterile milli-Q water, seeds were germinated on plates with solid 0.5x Murashige and Skoog (MS) medium containing 1% agar (without vitamins or sucrose). The medium was supplemented with kanamycin (100 μg/ml) when required to select transgenic plants. Plates were incubated in a climate-controlled growth chamber (Ibercex) at 26°C with a photoperiod of 14 h of white light (photon flux density of 50 μmol m^-2^ s^-1^) and 10 h of darkness. After 10-14 days, seedlings were transferred to soil and grown under standard greenhouse conditions (14 h light at 25 ± 1 °C and 10 h dark at 22 ± 1 °C). Plants used for root metabolic analysis were grown in a greenhouse with the same conditions but in a mixture of river sand (0.5-1 mm, Filcom BV) and a clay granulate (1:1) instead of soil for easier root collection. For the analysis of phenotypical traits influenced by SLs, fifteen plants were grown for 4 weeks under half strength Hoagland solution and then for 2 additional weeks under half strength Hoagland solution without PO ^-3^. *N. benthamiana* plants used for co-immunoprecipitation experiments were grown in the greenhouse under long day conditions at 24°C for 21 days.

### Sample collection and phenotypical analyses

Young leaf samples correspond to growing leaflets from the fifth and sixth true leaves and they were collected from soil-grown 4-week-old plants. Chlorophyll fluorescence measurements were carried out with a Handy FluorCam (Photon Systems Instruments). ɸPSII was measure at 30 PAR with an actinic light of 3 μmol m^-2^ s^-1^. Tomato fruit pericarp samples for isoprenoid quantification were collected at B+3 (three days after breaker). Roots, leaflets and pericarp samples were frozen in liquid nitrogen immediately after collection, freeze-dried and stored at -80 °C. For counting the days from breaker to orange stage, thirty fruits (n=30) from each genotype were chosen and their ripening was monitored *in planta*. For fruit weight determination, 100 ripe fruits from each genotype were collected and weighted one by one using a precision scale (Kern). Fruit volume was estimated in 10 pools of 10 fruits each by measuring the displaced water volume in a graduated cylinder.

### Constructs and tomato transformation

For co-immunoprecipitation experiments, full-length cDNAs encoding SlG1 and PSY3 proteins without their stop codons were amplified from root cDNA using the Phusion High-fidelity DNA polymerase (ThermoFisher). Next, the amplicons were introduced via BP clonase into pDONR207 entry plasmid using Gateway (GW) technology (Invitrogen). Full-length sequences were then subcloned through an LR reaction into pGWB414 and pGWB420 plasmids as previously reported (Barja et al., 2021). For CRISPR-Cas9-mediated disruption of *SlG1*, two single guide RNAs (sgRNA) (Figure S1-2) were designed using the online tool CRISPR-P 2.0 (Liu et al., 2017). Cloning of the CRISPR-Cas9 constructs was carried out as previously described (Barja et al., 2021) using primers listed in Table S1. As a result, a single final binary plasmid harboring the Cas9 sequence, the NPTII gene providing kanamycin resistance, and the sgRNAs was obtained and named pDE-Cas9-SlG1 (Table S2). All constructs were confirmed by restriction mapping and DNA sequencing. *Agrobacterium tumefaciens* GV3101 strain was used to stably transform tomato MicroTom cotyledons with pDE-Cas9-SlG1 as described (Ezquerro et al., 2022). *In vitro* regenerated T1 lines were identified based on kanamycin resistance (100 μg/ml) PCR genotyping and sequencing (Table S1). Homozygous T2 lines lacking Cas9 were obtained after segregation and stable T3 offspring was used for next experiments.

### Metabolite and gene expression analyses

Plastidial isoprenoids (carotenoids, chlorophylls, tocopherols), hormones (ABA, SL) and volatile organic compounds was carried out as described (Methods S1). Gene co-expression network**s** and RT-qPCR analyses were performed as described (Methods S2).

### *P. syringae* infection of tomato plants

*Pseudomonas syringae pv. tomato DC3000* (*Pst*) strain were used for tomato infection as previously reported (López-Gresa et al., 2018). Briefly, bacteria were grown during 48 h at 28°C in LB agar medium with rifampicin (10 mg/mL) and kanamicin (0.5 mg/mL). When colonies appeared, they were transferred to King’s B liquid medium supplemented with antibiotics and grown overnight at 28°C. Next, bacteria were centrifugated at 3000 x*g* for 15 min and resuspended in 10mM MgCl_2_ at a final optical density of 0.1 for further infection. Inoculation with bacteria was carried out in 4-week-old MicroTom plants without flowers by immersion. Plants were dipped into the bacterial suspension containing 0.05% Silwet L-77 for 30 seconds and left 24 hours for subsequent sample collection.

### Co-immunoprecipitation assays

Co-immunoprecipitation experiments were carried out as described (Barja & Rodríguez-Concepción, 2020; Barja et al., 2021) (Methods S3).

## Results and discussion

### Generation of CRISPR lines defective in SlG1

The approach followed to create *SlG1*-defective mutants by CRISPR-Cas9 was very similar to the one followed to generate *slg2* and *slg3* mutants (Barja et al., 2021). Briefly, we designed two single guide RNAs (sgRNA) (see Table S1 and S2 for primer and construct details) with CRISPR-P 2.0 (Liu et al., 2017) to create a small deletion that would disrupt the intronless *SlG1* gene (Figure 1). After transformation of tomato MicroTom plants and genotyping, two independent mutant alleles without Cas9 were selected and named *slg1-1* and *slg1-2* (Figure 1b).

The *slg1-1* allele has a deletion that causes a frameshift and a premature translation stop codon (Figure 1b and Figures S1-S2). The resulting protein lacks the C-terminal region containing the second Asp-rich motif (SARM, essential for prenyl-transferase function) and it is smaller than the wild-type (WT) enzyme (226 aa instead of 365 aa). A longer deletion in the *slg1-2* allele maintains the open reading frame and produces a 253 aa chimeric protein that lacks a fragment of the WT enzyme containing the SARM (Figure 1b and Figures S1-S2). Similar mutations lacking the C-terminal part of the protein and the SARM were previously shown to result in complete loss of GGPPS activity in *slg2* mutants (Barja et al., 2021). Therefore, we considered these two alleles as knock-out mutants and selected them for the rest of experiments.

### Loss of SlG1 does not impair the production of photosynthesis-related isoprenoids in tomato leaves

*SlG1* is expressed at low levels in all tomato plant tissues, including leaves (Figure S3). To investigate possible roles of SlG1 in leaves, we first analyzed the levels of GGPP-derived plastidial isoprenoids in *slg1* lines under normal growth conditions (Figure 2). Lines lacking SlG2 (*slg2-1*) or SlG3 (*slg3-1*) were grown together with the SlG1-defective mutants and WT controls for comparison. Young and mature leaves of *slg1-1* and *slg1-2* plants appeared very similar to those from *slg2* and WT plants (Figure 2a). By contrast, young emerging leaves from *slg3* plants showed a paler green color as previously reported (Barja et al., 2021). The color phenotype of young leaves correlated with their photosynthetic pigment (carotenoids and chlorophylls) content (Figure 2b) and their photosynthetic activity estimated as effective quantum yield of photosystem II (ɸPSII) (Figure 2c), which were only reduced in the *slg3* mutant. Tocopherol levels, by contrast, were similar in all the lines, although a trend towards lower levels was detected in young leaves of the *slg3* mutant, as previously reported (Barja et al., 2021). The described results are in agreement with our previous conclusion that *SlG3* is the main isoform in suppling GGPP for photosynthesis-related isoprenoids in leaves under normal growth conditions (Barja et al., 2021). A role for SlG2 in providing extra GGPP to support the production of these isoprenoids when needed, e.g., during deetiolation, was proposed in part based on gene expression data (Barja et al., 2021). Unlike *SlG2*, however, *SlG1* is poorly co-expressed with isoprenoid biosynthetic genes in leaves and it is not induced during seedling deetiolation (Barja et al., 2021), supporting the conclusion that SlG1 does not substantially contribute to GGPP production in chloroplasts.

**Figure 2.**
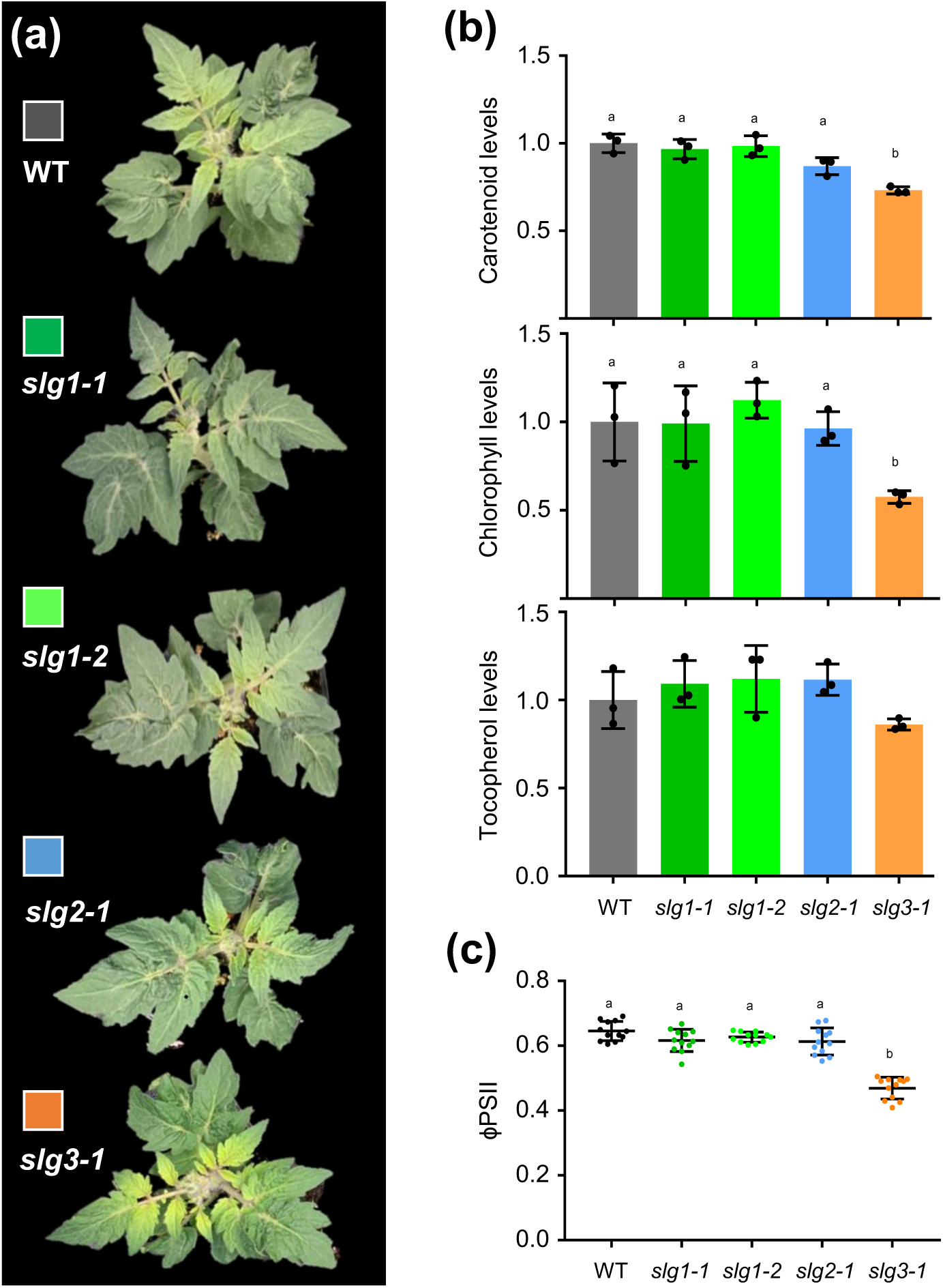
SlG1 does not contribute to GGPP-derived isoprenoid biosynthesis in tomato leaves. **(a)** Representative images of 4-week-old plants of WT and GGPPS-defective mutants. **(b)** Carotenoid, chlorophyll and tocopherol levels in young leaves from 4-week-old WT and mutant plants. Values are represented relative to WT levels and they correspond to mean and SD of n≥3 independent biological replicates. **(c)** Effective quantum yield of photosystem II (ɸPSII) in young leaves like those used in (b). Individual values (colored dots) and well as mean and SD are shown, and they correspond to four different areas from leaves of three different plants. In all plots, letters represent statistically significant differences (*P* < 0.05) among means according to post hoc Tukey’s tests run when one way ANOVA detected different means.

### SlG1 is not required for monoterpene synthesis in leaves

Leaves are mainly photosynthetic organs but they also contain cell types lacking chloroplasts. In particular, tomato leaves contain large amounts of glandular trichomes formed by non-photosynthetic cells that produce large amounts of volatile organic compounds (VOCs), including many of isoprenoid origin (Schuurink and Tissier, 2020). *SlG1* but also *SlG2* and *SlG3* are expressed in leaf trichomes (Zhou & Pichersky, 2020). The main groups of isoprenoid VOCs are MVA-derived sesquiterpenes (made from C15 FPP), MEP derived monoterpenes (made from C10 GPP or nerolidol diphosphate, NPP) and diterpenes (made from C20 GGPP). A role for SlG1 in the production of GGPP-derived diterpene VOCs was investigated following the observation that *SlG1* expression was induced by treatments that stimulated the production of such VOCs (Ament et al., 2006). Our data mining of the database Genevestigator found that *SlG1* expression is also upregulated in leaves after infection with the bacterium *Pseudomonas syringae pv. tomato* DC3000 (*Pst*), whereas *SlG2* and *SlG3* transcript levels remained unchanged (Figure S4). The most prominent isoprenoid VOCs produced upon *Pst* infection are monoterpenes such as linalool, limonene and α-terpineol (Lopez-Gresa et al., 2017; Zhou & Pichersky, 2020). Linalool is produced from GPP synthesized by homodimeric GPP synthases such as tomato GPPS (Solyc08g023470) or heterodimeric enzymes formed by a GGPPS subunit and the small subunit type I (SSU-I) protein (Solyc07g064660), whereas limonene and α-terpineol derive from NPP produced by the NPP synthase NDPS1/CPT1 (Solyc08g005680) (Schilmiller et al., 2009) (Figure 3a). Similar to *SlG1*, tomato *NDPS1*, *GPPS* and *SSU-I* genes are induced in *Pst*-infected leaves, with *SlG1, SSU-I* and *NDPS1* showing the strongest upregulation (Figure S4). SlG1/SSU-I heterodimers have been shown to mainly produce GPP *in vitro* (Zhou & Pichersky, 2020) and SlG2/SSU-I heterodimers have been found to produce GPP for monoterpene biosynthesis in tomato fruit (Hivert et al., 2020). Based on these data, we hypothesized that SlG1 might participate in the production of monoterpenes in leaves.

**Figure 3.**
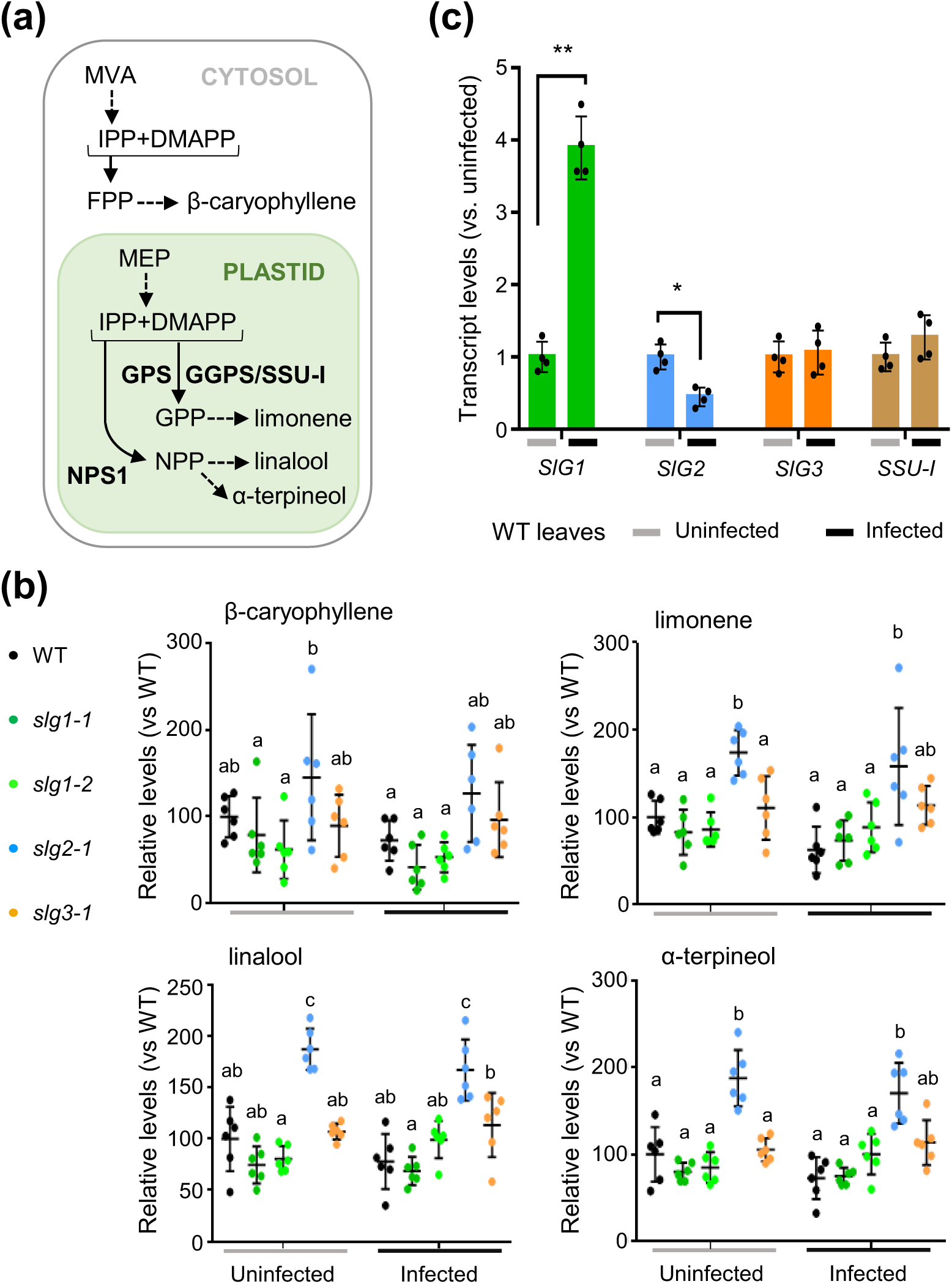
SlG1 is not required for monoterpene production in leaves. MicroTom WT and edited plants were infected with *Pseudomonas syringae pv. tomato* DC3000 and samples were collected 24 h later. **(a)** Biosynthetic pathways. **(b)** Levels of representative VOCs in infected and uninfected leaves of WT and GGPPS-defective mutants. Data are represented relative to the levels in uninfected WT samples (100%) and correspond to the mean ± SD of n=6 biological replicates. Letters represent statistically significant differences (one-way ANOVA followed by Tukey’s multiple comparisons test, P < 0.05). A schematic representation of the VOC biosynthesis pathway is shown in the left side. **(c)** RT-qPCR expression data of the indicated genes in WT leaves. Expression levels are shown relative to uninfected samples and they correspond to the mean ± SD of n=4 independent biological replicates. Asterisks indicate statistically significant differences among means between uninfected and infected samples (t-test: *, P < 0.05; **, P < 0.01).

To test this hypothesis, we infected WT and GGPPS-defective tomato plants with *Pst* and quantified the levels of several monoterpenes (and a sesquiterpene as a control) before and 24 h after the infection (Figure 3b). Strikingly, bacterial infection did not cause significant changes in monoterpene or sesquiterpene levels in WT plants or any of the mutants tested (Figure 3b) even though a clear upregulation of *SlG1* expression was observed (Figure 3c). Surprisingly, *SlG2* transcript levels decreased in *Pst*-infected WT samples (Figure 3c) while *slg2* mutants contained increased basal levels of all VOCs analyzed (Figure 3b). These results suggest that impairment of SlG2 activity results in higher production and/or accumulation of these VOCs, which are derived from IPP and DMAPP from different origins: the MVA pathway (via FPP) and the MEP pathway (via NPP or GPP). However, the levels of leaf isoprenoid products made from MEP-derived GGPP (carotenoids, chlorophylls, tocopherols) were similar in *slg2* and WT lines (Figure 2b) (Barja et al., 2021). We speculate that loss of SlG2 function in trichomes (the VOC-producing cells in the leaf) might cause the diversion of SlG2 substrates (i.e., IPP and DMAPP) to the production of VOCs. Alternatively, a metabolic imbalance or a defense response in *slg2* plants that cannot be rescued by SlG1 or SlG3 might eventually lead to VOC overproduction.

Unlike the results from Genevestigator RNA-seq data (Figure S4), in our experiment we did not detect any changes in the levels of *SSU-I* transcripts (Figure 3c). This Genevestigator differential expression profile might be a consequence of different genetic backgrounds used in the experiments. In agreement with this hypothesis, *SSU-I* transcript levels are very low in cultivated tomatoes (*S. lycopersicum*) but are abundant in the fruits of wild relatives (*S. pimpinellifolium* and *S. cheesmaniae*), which produce substantially more monoterpenes (Hivert et al., 2020). Whereas MicroTom has been broadly used for infection experiments with some bacteria and fungi (Costa et al., 2021; Deganello et al., 2014; Nakahara et al., 2016), previous experiments using *Pst* found no wilt symptoms or bacterial growth eight days after infection (Takahashi et al., 2005), strongly suggesting that *Pst* is just an opportunistic MicroTom pathogen. This suggests that tomato MicroTom is not the best genotype to investigate the molecular and metabolic changes triggered by *Pst* infection. However, the similar levels of monoterpenes found in non-infected leaves of WT and SlG1-defective alleles argues against a role of SlG1 in the production of these plastidial isoprenoids in leaves.

### SlG1 is dispensable for carotenoid biosynthesis in fruit

Carotenoids are synthesized at very high rates during tomato fruit ripening, contributing together with the degradation of chlorophylls to progressively change the fruit color from green at the mature green (MG) stage to orange (O) and eventually red at the ripe (R) stage. The first visual symptoms of color change define the breaker (B) stage. *SlG1* is expressed at very low levels during fruit ripening (Figure S3), when GGPP produced by SlG3 and upregulated levels of *SlG2* support carotenoid overproduction (Barja et al., 2021). Reduced activity of SlG2 and SlG3 results in lower levels of lycopene (the main carotenoid accumulated during ripening) in R fruit collected at 10 days after the B stage (B+10) but only SlG3-defective lines showed significantly decreased levels of total carotenoids at this stage and none of the mutants showed differences with the WT in MG fruit (Barja et al., 2021). When we measured carotenoid levels in B+3 fruits (*i.e.,* between O and R stages), WT levels of total carotenoids were found in *slg1* and *slg2* lines, whereas *slg3* fruits showed significantly lower levels (Figure 4a). In particular, phytoene and lycopene contents were strongly reduced in *slg3* fruits, whereas β-carotene levels were similar to those of WT controls (Figure 4a). The absence of significant differences in carotenoid levels among WT, *slg1* and *slg2* fruit at the B+3 stage (Figure 4a) suggests that SlG3 is the main GGPP provider in the early stages of fruit ripening. As ripening advances, up-regulation of SlG2 contributes with extra GGPP. The WT phenotype of SlG1-deficient mutant fruits together with the lack of gene expression changes during ripening support the conclusion that this isoform does not contribute to GGPP for carotenoid production in fruit.

**Figure 4.**
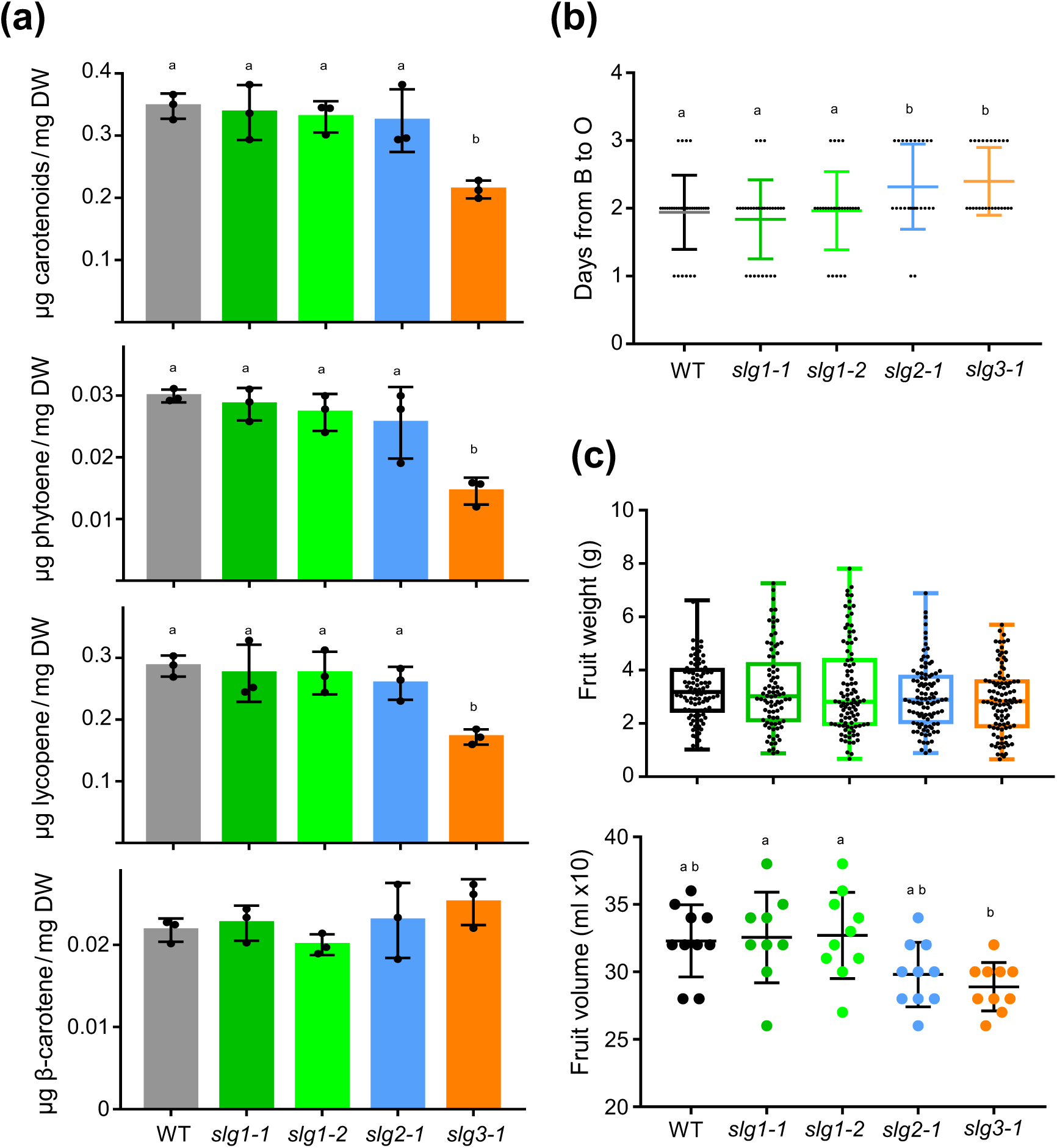
SlG1 is dispensable for carotenoid biosynthesis in fruits. **(a)** Levels of total and individual carotenoids (phytoene, lycopene and β-carotene) in tomato fruits collected from the plant 3 days after the breaker stage (B+3) fruits. Values represent mean and SD of n=3 independent biological replicates**. (b)** Fruit ripening rate estimated as the number of days from B to O stages in the plant. Black dots indicate individual values and colored lines represent the mean and the SD. **(c)** Weight and volume of fully ripe (R) fruits of the indicated genotypes. In the weight boxplot, the lower and upper boundary of the boxes indicate the 25th and 75th percentile, respectively; the line inside the boxes represents the median; dots mark individual data values; and whiskers above and below the boxes indicate the maximum and minimum values. In the volume dotplot, central line represents the mean and whiskers represent SD. In all cases, letters represent statistically significant differences (P < 0.05) among means according to post hoc Tukey’s tests run when one way ANOVA detected different means.

Lower levels of the carotenoid-derived hormone ABA were measured in the fruit pericarp of tomato mutants lacking SlG2 and particularly SlG3, eventually contributing to a delay in ripening (Barja et al., 2021). Consistently, the number of days that B fruits needed to reach the O stage was higher in *slg2* and *slg3* lines compared to WT controls (Figure 4b). ABA has also been shown to promote fruit growth (McQuinn et al., 2020; Zhang et al., 2009; Ezquerro et al. 2022). Reduced ABA contents in *slg2* and *slg3* fruit pericarp (Barja et al., 2021) actually correlate with a reduced fruit volume of ripe fruit, although this was only statistically significant for *slg3* fruits and it did not affect fruit weight (Figure 4c). Again, *slg1* fruits were undistinguishable from WT controls (Figure 4c). The observation that losing SlG1 activity does not impact any of the fruit phenotypes tested strongly supports the conclusion that this isoform is dispensable for the production of GGPP for carotenoids and related metabolites during fruit ripening.

### Gene co-expression analysis suggests a major role for SlG1 in roots

In our previous work we demonstrated that *SlG2* and, to a lower extent, *SlG3* expression were highly connected to the expression of plastidial isoprenoid biosynthetic genes in leaf tissue. In fruits, *SlG3* had a higher connection than *SlG2* to genes from these metabolic pathways (Barja et al., 2021). The correlation of *SlG1* with other plastidial isoprenoid genes was very poor in leaves as well as in fruits. Other gene expression data suggested that *SlG1* might function in roots to produce SL or/and AM-related apocarotenoids (Stauder et al., 2018; Barja et al. 2021). To provide further evidence for this hypothesis, we performed a gene co-expression network (GCN) analysis in roots. We used publicly available data for plant comparative genomics (PLAZA 4.0 Phytozome) to look for tomato homologues of genes for plastidial isoprenoid biosynthesis and related pathways. We obtained the expression data of such tomato homologs from recently published tomato RNA-seq data in root tissue (Wang et al., 2021) and calculated their expression correlation with *SlG1*, *SlG2* and *SlG3* expression using pairwise Pearson correlations as previously reported (Wang et al., 2022). Results are shown in Figure 5 and correlations and gene details are listed in Table S3. Opposite to that observed in leaf tissue, *SlG2* displays a low correlation with other isoprenoid biosynthetic genes in roots (Figure 5). *SlG3* shows a medium connectivity to many of the selected genes, probably because it is the isoform providing GGPP for housekeeping functions (Barja et al., 2021). Intriguingly, *SlG1* showed high correlation with many genes of the MEP and the carotenoid pathways as well as with most of the genes involved in SL biosynthesis, while it showed limited connectivity with genes converting carotenoids into ABA (Figure 5). These results suggest that gene expression is coordinated such that SlG1-derived root carotenoids are channeled into the production of SLs but not ABA. *SlG1* connectivity was also very high with GA biosynthetic genes. GAs are GGPP-derived phytohormones that appear to act together with SLs during the P-starvation response (Wang et al., 2021) and root mycorrhization (Nouri et al., 2021; Ruiz-Lozano et al., 2016), promoting root growth to help establishment of AM symbiosis. A tight correlation between SLs and the expression of GA-related genes has been reported (Wang et al., 2021), suggesting that the observed co-expression of *SlG1* with GA biosynthetic genes might be a secondary effect.

**Figure 5.**
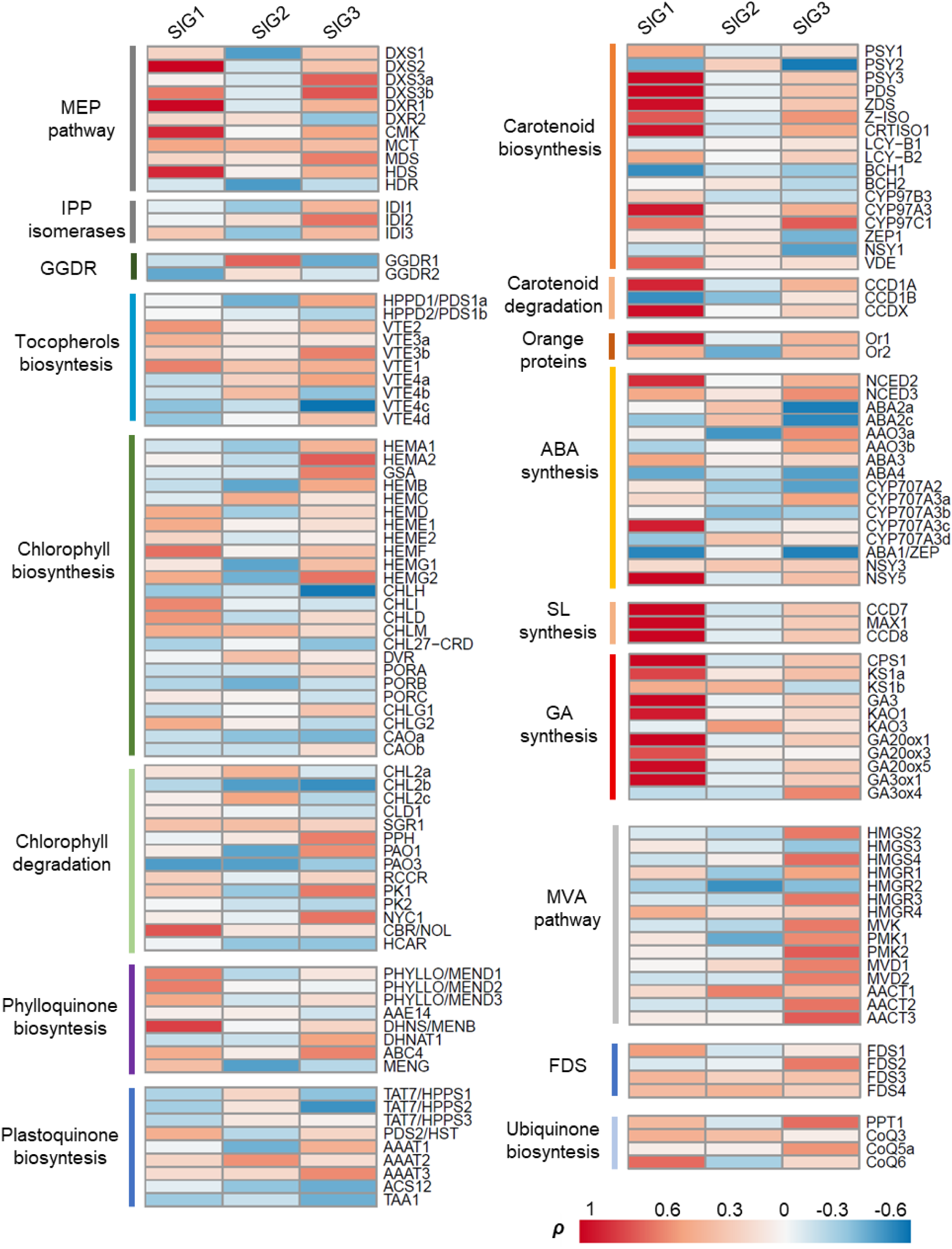
*SlG1* expression is highly connected with carotenoid and SL biosynthesis in roots. Heatmap represents pairwise Pearson correlations (ρ) between the expression of genes encoding GGPPS isoforms and those for the indicated enzymes from pathways upstream and downstream of GGPP. Gene details, abbreviations, accessions, and data correlations are listed in Table S3.

### SlG1 specifically interacts with PSY3

Gene expression data and our GCN analysis (Figure 5), support the conclusion that SlG1 might function in coordination with PSY3 to produce SL and likely other AM-associated apocarotenoids (but not ABA) in roots (Barja et al., 2021; Stauder et al., 2018). GGPPS proteins have been shown to physically interact with PSY enzymes in different plant species (Ruiz-Sola et al., 2016b; Wang et al., 2018; Camagna et al., 2019; Barja et al., 2021). In tomato, co-immunoprecipitation experiments in *Nicotiana benthamiana* leaves showed direct interaction of SlG2 with PSY1 and PSY2, while no interaction with these PSY isoforms was found for SlG3 even though the latter was shown to homodimerize and heterodimerize with SlG2 using the same experimental design (Barja et al., 2021). To extend the tomato GGPPS-PSY interaction map with SlG1, we performed co-immunoprecipitation assays *in planta* using Myc-tagged GGPPS and HA-tagged PSY proteins (Figure 6). Tagged proteins were transiently expressed in *N. benthamiana* leaves by agroinfiltration and then protein extracts were used to confirm the presence of the recombinant enzymes (Figure 6a) and for co-immunoprecipitation with anti-Myc antibodies followed by immunoblot analysis with both anti-Myc and anti-HA antibodies (Figure 6b). A Myc-tagged phosphoribulokinase protein from Arabidopsis (PRK-Myc) was used as a negative control (Barja et al., 2021). Additionally, a SlG1-HA construct used together with SlG1-Myc confirmed that SlG1 forms homodimers and that the Myc-tagged version allowed to co-immunoprecipitate protein partners (Figure S5). When combined with PSY isoforms, SlG1 was found to only co-immunoprecipitate with PSY3, whereas SlG2 and SlG3 were unable to interact with this particular isoform (Figure 6b).

**Figure 6.**
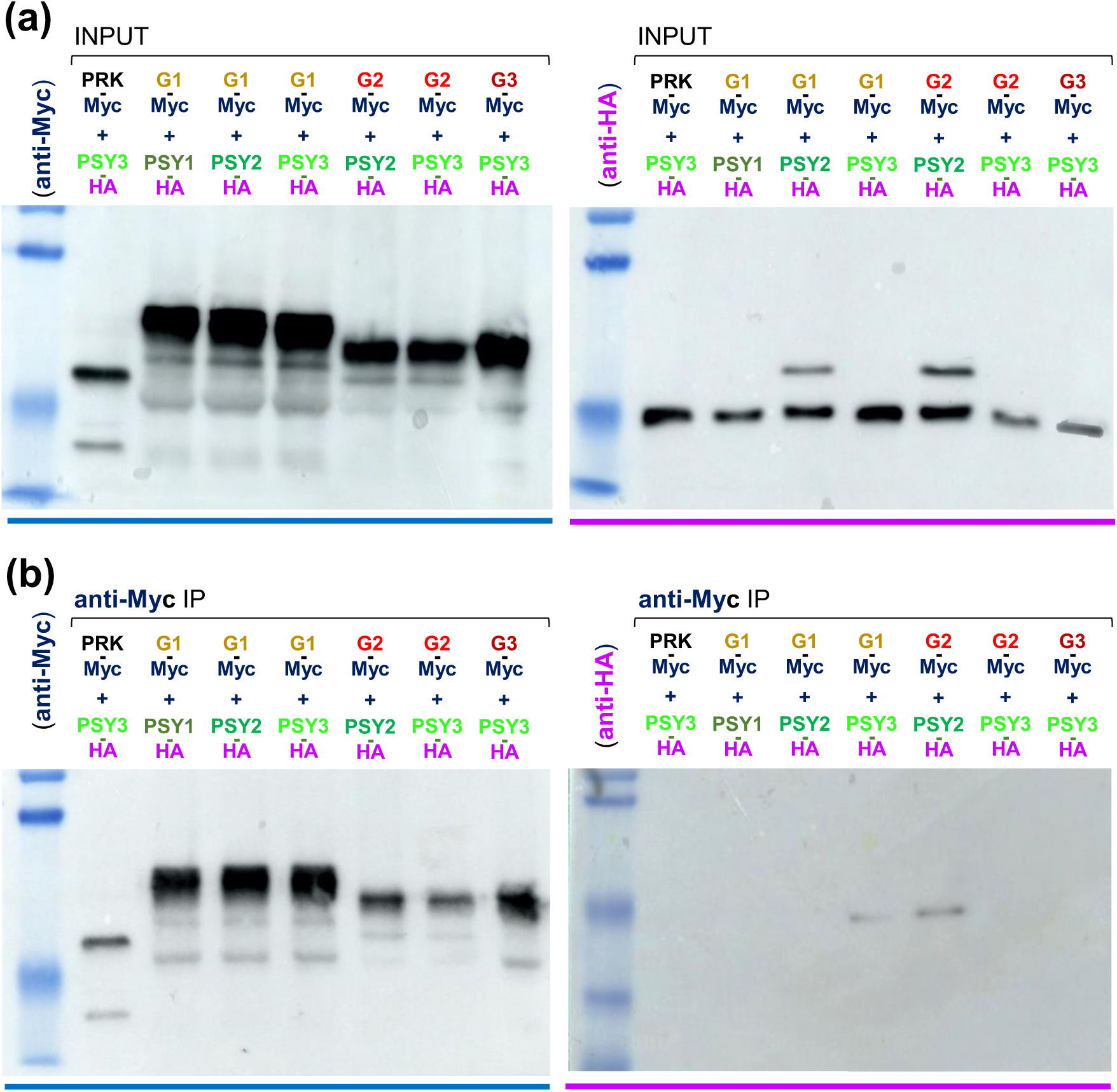
SlG1 specifically interacts with PSY3 *in planta*. *N. benthamiana* leaves were co-agroinfiltrated with constructs encoding the indicated proteins tagged with C-terminal Myc or HA epitopes. **(a)** Immunoblot analysis of crude extracts (INPUT) with anti-Myc (dark blue) and anti-HA (purple) antibodies to confirm successful protein production. **(b)** Immunoblot analysis of extracts after immunoprecipitation (IP) with anti-Myc. The same samples were used for immunodetection with anti-Myc (to confirm successful IP) or anti-HA (to identify co-immunoprecipitated partners). Predicted protein molecular weights (KDa): PRK-Myc, 52.1; SlG1-Myc, 56.9; SlG2-Myc, 55.3; SlG3-Myc, 55.5; PSY1-HA, 50.7; PSY2-HA, 51.0; PSY3-HA, 50.2.

It has been suggested that PSY enzymes cannot access freely diffusible plastidial GGPP because of their specific plastid location attached to membranes (Camagna et al., 2019), making interaction among GGPPS and PSY enzymes necessary for GGPP channeling into the carotenoid pathway. Strikingly, the tomato housekeeping isoform SlG3 is unable to directly interact with any of the PSY isoforms present in tomato (Figure 6b) (Barja et al., 2021). However, there are several possibilities for an indirect interaction of SlG3 with PSY enzymes (Barja & Rodriguez-Concepcion, 2021). Heterodimerization of SlG2 and SlG3 might allow interaction of SlG3 with PSY1 or PSY2 via SlG2 (Barja et al., 2021), whereas possible heterodimerization of SlG1 and SlG3 might allow interaction with PSY3 (Figure 6b). Another mechanisms that could potentially facilitate the GGPP - PSY interaction could be interaction with the small subunit type II (SSU-II) protein (Solyc09g008920), a catalytically inactive polypeptide shown to interact with different GGPPS enzymes to improve their GGPP production (Wang et al., 2018; Zhou et al., 2017; Zhou & Pichersky, 2020) but also to stimulate the interaction with PSYs (Wang et al., 2018). While the described interactions potentially allow any of the three tomato GGPPS isoforms to form a complex with any of the PSY isoforms, it is expected that direct interactions (SlG1-PSY3, SlG2-PSY1 and SlG2-PSY2) would be most efficient to convert GGPP into phytoene. In the case of SlG1 and PSY3, this interaction is strengthened by the coordinated expression profiles of the corresponding genes in roots upon phosphate starvation and mycorrhization with AM fungi (Figure 5) (Barja et al., 2021; Stauder et al., 2018) and this strongly supports the conclusion that these isoforms share the same functional role(s).

### SL synthesis is impaired in *slg1* and *psy1* mutants

To experimentally confirm the role of the different GGPPS and PSY isoforms in the production of carotenoid precursors for SL or/and ABA biosynthesis in roots, we analyzed the levels of these metabolites in WT plants and mutant lines defective in GGPPS and PSY isoforms. In the latter case, only CRISPR-Cas9 lines lacking PSY1 (*psy1-2*) and PSY2 (*psy2-1*) were used (Ezquerro et al., 2022), as PSY3-defective lines are not available. As a control, we used a tomato MicroTom line with a silenced *CCD7* gene (Solyc01g090660), encoding the SL biosynthetic enzyme carotenoid-cleavage dioxygenase 7 (Pino et al., 2022). This line, herein named *ccd7*, shows a highly branched phenotype consistent with expectedly low SL production (Pino et al., 2022). We grew the plants in half-strength Hoagland solution either containing normal phosphate (+P) or without phosphate (-P) to induce SL synthesis and measured carotenoid and ABA levels in root tissues and SL levels in root exudates (Figure 7). Carotenoid levels measured in roots were very low in all genotypes both under +P and -P conditions (Figure 7a). Only the *ccd7* line displayed statistically significant changes between conditions, as carotenoid levels increased under phosphate starvation (Figure 7a).

**Figure 7.**
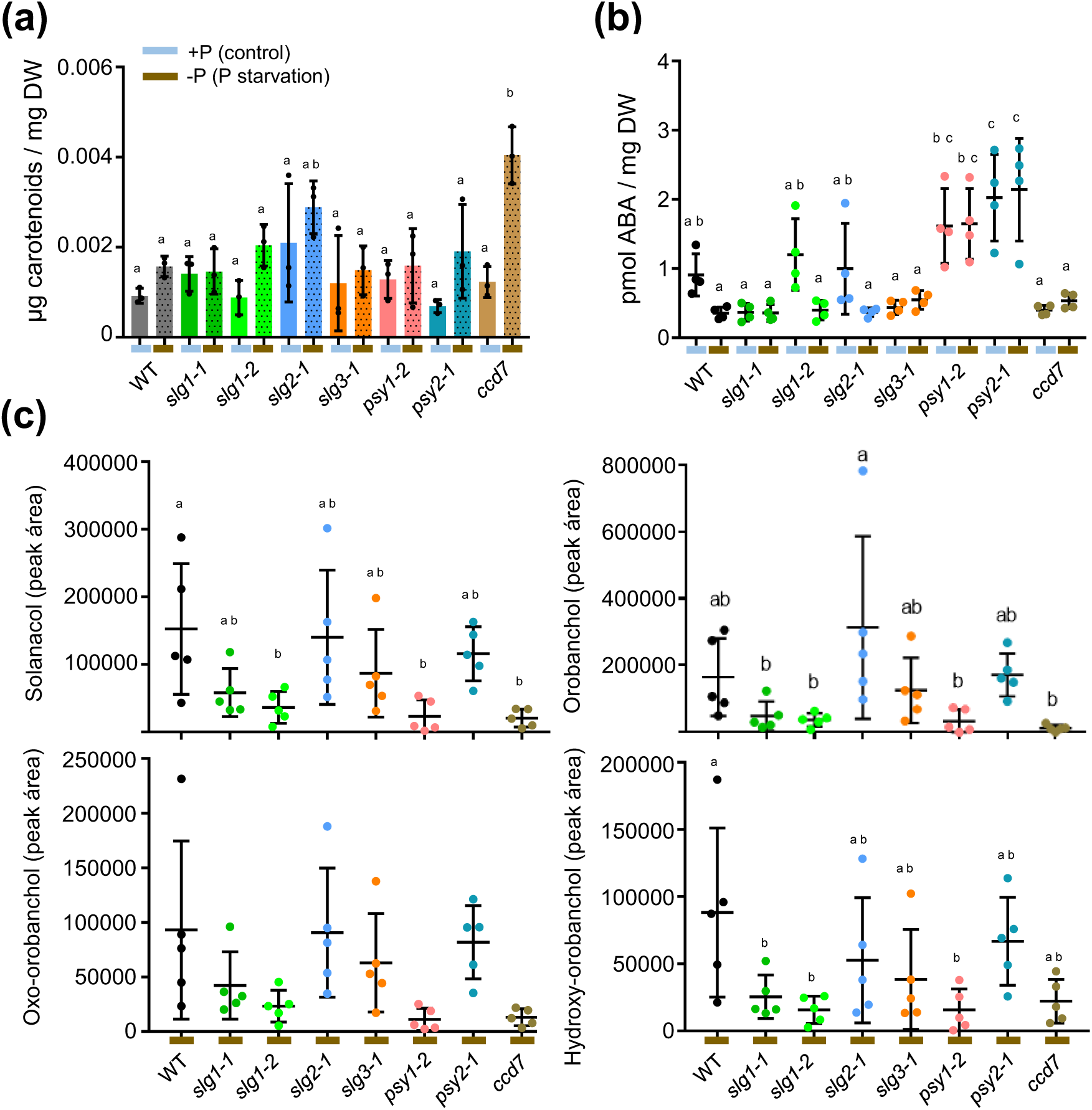
SlG1 is involved in root SL production. Plants of the indicated genotypes were grown in half-strength Hoagland solution with normal phosphate (+P) or under phosphate starvation (-P) conditions. Samples of root tissues or exudates were collected for metabolite analyses. **(a)** Carotenoid levels in root tissues. Values correspond to the mean and SD of n≥3 independent biological replicates. **(b)** ABA levels in root tissues. Values correspond to mean and SD of n≥4 independent biological replicates. **(c)** Levels of individual SLs in root exudates. Values represent the mean and SD of n≥5 independent biological replicates. In dotplots, inner line is the mean and whiskers represent SD. In all cases, letters represent statistically significant differences (p<0.05) among means according to posthoc Tukey’s tests run when one-way ANOVA detected different means.

Next, we measured the levels of the carotenoid-derived phytohormone ABA in the same samples used for carotenoid quantification. Consistent with our GCN data showing no correlation of ABA biosynthetic genes with any of the tomato genes for GGPPS isoforms (Figure 5), none of the GGPPS-defective mutants presented statistically significant differences in root ABA levels in either +P or -P conditions compared to WT controls (Figure 7b). Interestingly, ABA levels were higher in roots of *psy1-2* and *psy2-1* plants grown under +P and –P conditions (Figure 7b). Analysis of gene expression levels by qRT-PCR showed that loss of PSY1 and PSY2 activities did not substantially increase the expression of any remaining PSY-encoding gene or any GGPPS-encoding gene in roots (Figure 8). We therefore conclude that other genes involved in ABA synthesis might be upregulated in *psy1* and *psy2* roots. Alternatively, loss of PSY1 or PSY2 in roots might cause a metabolic imbalance that cannot be rescued by the remaining PSY isoforms and eventually leads to ABA overproduction.

**Figure 8.**
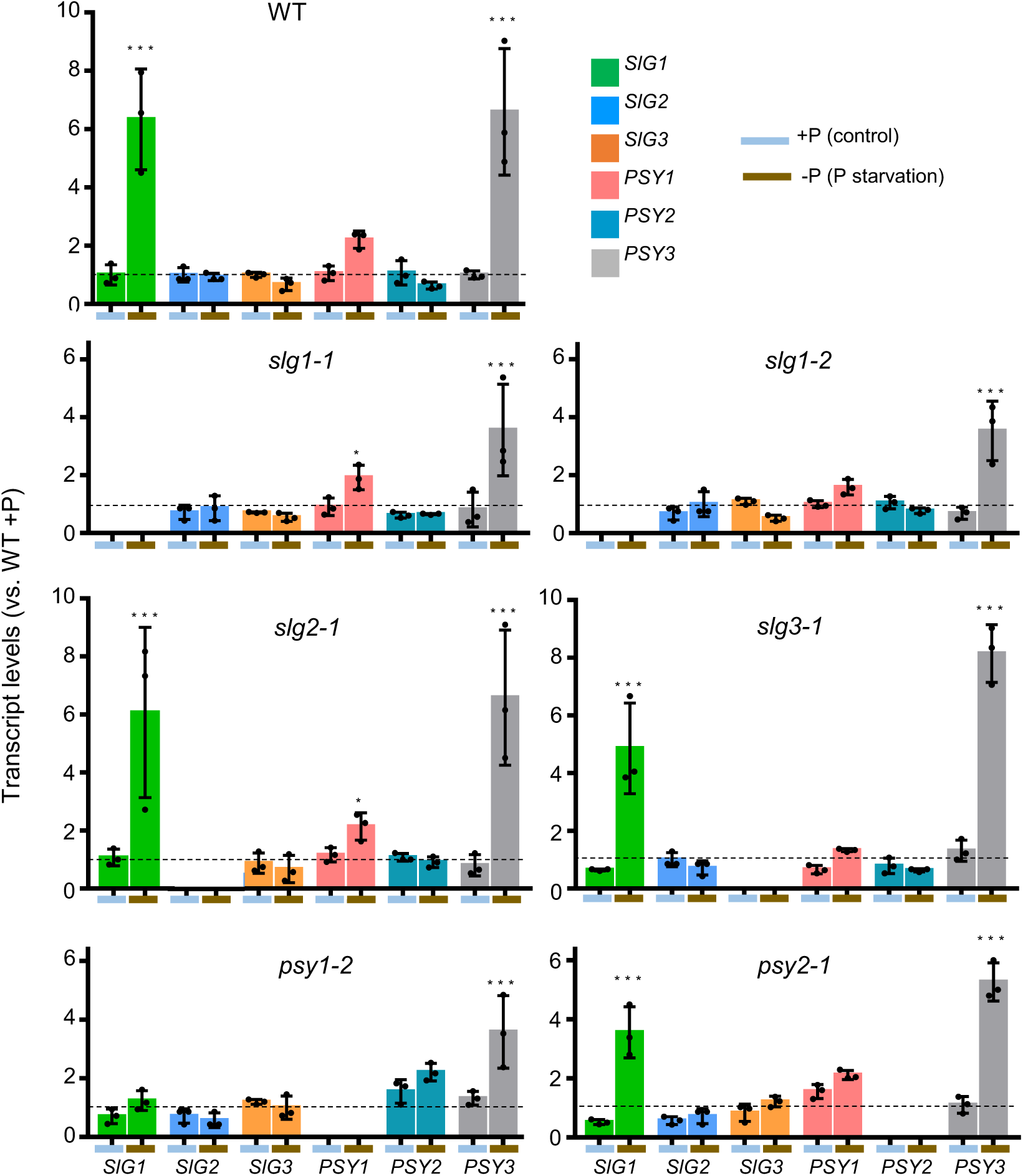
Genes encoding GGPPS and PSY paralogues show differential responses to phosphate starvation in roots. RNA samples from roots collected from the plants described in Figure 7 were used for RT-qPCR experiments. Transcript levels were normalized using the tomato *ACT4* gene and they are shown relative to those in control (+P) WT samples (dotted line). The scale is the same in all plots to facilitate comparisons. Mean and SD of n=3 independent biological replicates are shown. Asterisks indicate statistically significant differences between conditions (+P vs -P) for each gene in each genotype according to one-way ANOVA with Dunnett’s multiple comparisons test: *, *P* < 0.05; **, *P* < 0.01; ***, *P* < 0.001).

The levels of several SLs were next measured in root exudates from plants grown under phosphate starvation for seven days (Figure 7c). All SL measured (except for oxo-orobanchol which also displays a trend towards lower levels) were significantly reduced in both *slg1* mutant alleles compared to WT controls, confirming a major role for the SlG1 isoform in producing GGPP precursors for these carotenoid-derived hormones. Exudates from *slg2-1* and *slg3-1* roots contained WT levels of SLs (Figure 7c). However, the observation that none of the two null *slg1* mutant alleles showed a complete absence of SLs suggest that SlG2 or/and SlG3 can also contribute to SL production, at least when SlG1 activity is absent. Because a role for PSY3 in root SL production has been demonstrated in the legume *Medicago truncatula* and suggested in tomato (Stauder et al., 2018), we expected WT levels of SLs in mutants defective in PSY1 and PSY2, which harbor a functional PSY3 enzyme. However, *psy1-2* displayed a very similar reduction in SL levels to that detected in *slg1*, suggesting that PSY1 may also have a role in SL synthesis in tomato roots (Figure 7c). In agreement, *PSY1* basal expression levels in roots are higher than those of *PSY3* (Barja et al., 2021; Fantini et al. 2013) and they increase in mycorrhized roots compared to non-mycorrhized controls (Barja et al., 2021; Stauder et al., 2018). Strikingly, in PSY1-defective roots SlG1 expression was not upregulated under phosphate starvation (Figure 8), suggesting that this might be the main cause of the low SL levels produced by the *psy1-2* mutant (Figure 7c). This result also reinforces the conclusion that SlG1 has a central role in SL production in roots. Furthermore, upregulation of *PSY3* expression under phosphate starvation was also reduced (but not impaired) in *psy1-2* roots, similar to that observed in SlG1-defective roots (Figure 8). This result suggests that SLs might feedback promote *PSY3* expression. Alternatively, the absence of SlG1 (in *slg1* mutants) or the failure to upregulate its levels under phosphate starvation (in *psy1* mutants) might be the reason why *PSY3* expression also becomes less responsive to phosphate starvation. As SlG1 and PSY3 isoforms physically interact (Figure 6), it is not surprising that their transcription is coordinated. In any case, the data strongly support a central role for SlG1 and PSY3 in root SL production.

### SL reduction in *slg1* and *psy1* roots does not affect aerial plant architecture

SLs are plant phytohormones that regulate developmental processes in roots, shoot and leaves. They promote root hair elongation, lateral root outgrowth and primary root growth and inhibit adventitious root formation (Al-Babili & Bouwmeester, 2015; Matthys et al., 2016; Ruyter-Spira et al., 2011; Kohlen et al., 2012). In shoots, they promote secondary growth and inhibit auxiliary bud branching (Gomez-Roldan et al., 2008; Ruyter-Spira et al., 2013), and in leaves they promote leaf senescence together with other plant hormones (Ueda & Kusaba, 2015; Yamada & Umehara, 2015). While increased branching is one of the most conspicuous phenotypes caused by reduced SL levels, a visual inspection could not detect any obvious branching phenotype in greenhouse-grown plants of any of our CRISPR-Cas9-edited lines.

To obtain quantitative data, we measured several phenotypic parameters related with SL-regulated plant growth in these lines grown together with WT controls and SL-deficient *ccd7* plants (Pino et al., 2022; Vogel et al., 2010). Measurements were performed on fifteen plants per genotype grown under +P for four weeks and then transferred to -P for two more weeks to promote SL biosynthesis (Ruyter-Spira et al., 2013) (Figure 9). Dry root weight was reduced in SL-deficient *slg1-1*, *slg1-2*, *psy1-2* and *cdd7* lines compared to WT controls and mutants with normal SL production, i.e., *slg2-1*, *slg3-1* and *psy2-1* plants (Figure 9a). These data confirm that SLs have a role in promoting root growth (Al-Babili & Bouwmeester, 2015; Ruyter-Spira et al., 2013). Next, we measured the height of the plants and observed that *ccd7* plants were smaller than WT plants (Figure 9b). Surprisingly, the SL-deficient mutants (*slg1-1*, *slg1-2* and *psy1-2*) showed a very similar size as WT plants, whereas *slg2-1* and *slg3-1*, which produced normal levels of SLs in roots, were smaller (Figure 9b). The reduced size of *slg2-1* and *slg3-1* plants might be derived from their metabolic imbalances resulting from suboptimal photosynthesis (Barja et al., 2021). SL-synthesis mutants typically are dwarfs with increased numbers of lateral branches (Gomez-Roldan et al., 2008; Yamada et al., 2014). Indeed, *ccd7* plants not only were smaller (Figure 9b) but also displayed a higher number of lateral branches compared to WT controls in our experimental conditions (Figure 9c), consistent with previous reports by other groups (Pino et al., 2022; Vogel et al., 2010). In agreement with our preliminar visual observations, none of the mutants defective in GGPPS or PSY isoforms showed a branching phenotype (Figure 9c). The results indicate that only *ccd7* plants display a SL deficiency shoot phenotype, probably because constitutive silencing of the *CCD7* gene affects SL production in all plant tissues. By contrast, loss of SlG1 appears to only affect SL production in roots, impacting root growth but not shoot growth or branching. The similar phenotypes of *slg1-1*, *slg1-2* and *psy1-2* plants further support the conclusion that the reduced SL production by PSY1-defective roots is due to a block in *SlG1* upregulation rather than caused by decreased PSY1 activity. It is commonly accepted that SLs are synthetized in plant roots and from there transported to aerial parts, where they regulate plant growth and shoot branching (Al-Babili & Bouwmeester, 2015; Gomez-Roldan et al., 2008; Ruyter-Spira et al., 2013). Nevertheless, grafting studies in tomato and pea have shown that the SL biosynthetic machinery is active in stems and can produce SLs that are transported via the xylem to shoots and leaves (Beveridge et al., 2009; Visentin et al., 2016; Xie et al., 2010). As the expression levels of *SlG1* in aerial parts under normal conditions are very low (about 40-fold lower than *SlG2* and 50-fold lower than *SlG3*) (Figure S3) (Barja et al., 2021; Stauder et al., 2018), we propose that SlG1 is exclusively involved in providing GGPP for SL synthesis in roots. In shoots, much higher levels of carotenoids (derived from GGPP made by SlG2 and/or SlG3) may supply enough precursors to produce SLs supporting normal aboveground growth and development.

**Figure 9.**
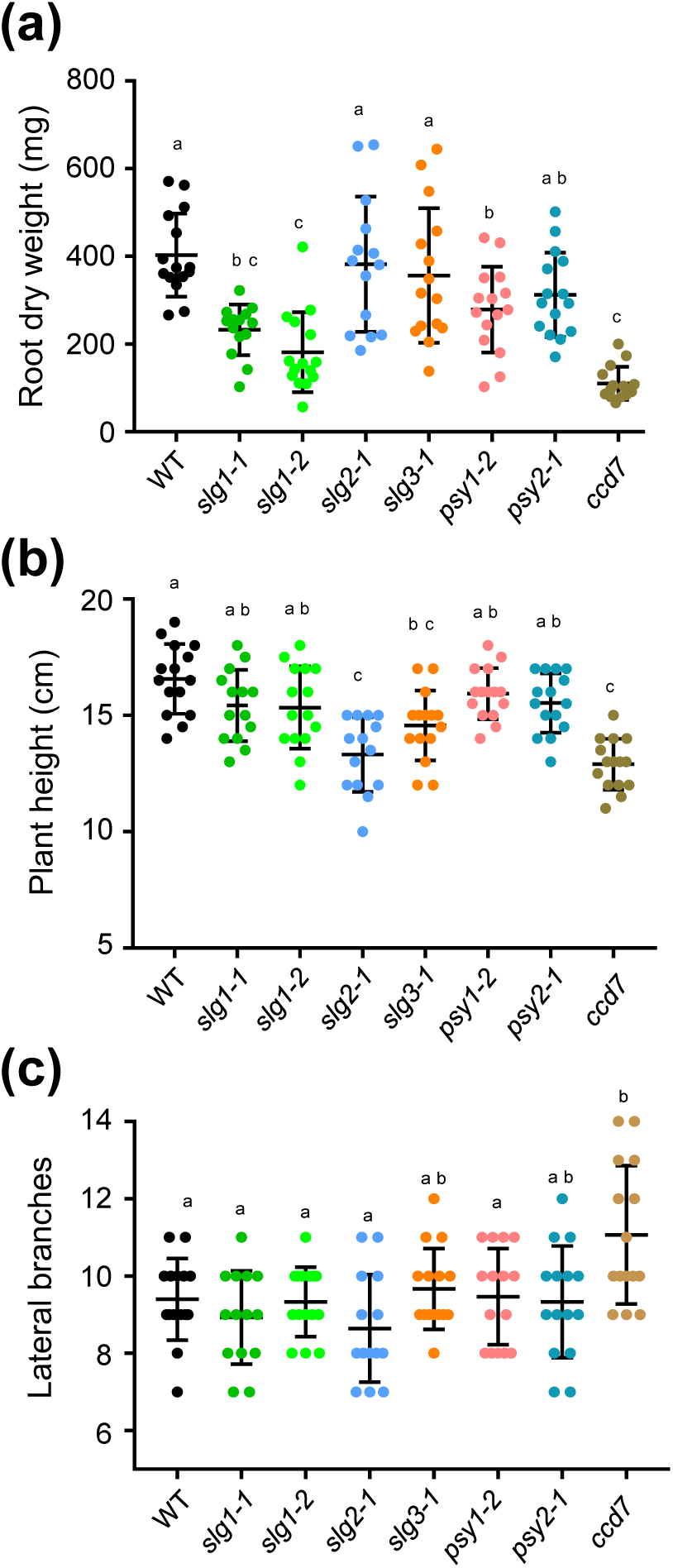
Defective SL synthesis in *slg1* plants does not impact shoot growth or branching. Measurements were performed using plants of the indicated genotypes grown under +P for 4 weeks and then transferred to -P for 2 more week to promote SL synthesis. **(a)** Weight of entire roots after freeze-drying. **(b)** Plant height from the root-stem transition to the apical meristem bud. **(c)** Number of lateral branches arising from the main stem. In all plots, dots indicate individual values, whiskers indicate mean and SD, and letters represent statistically significant differences (*P* < 0.05) among means according to posthoc Tukey’s tests run when one-way ANOVA detected different means.

## Concluding remarks

In summary, extensive characterization of tomato lines defective in the SlG1 isoform only found a clear phenotype in the root, consistent with the high connectivity of *SlG1* to other isoprenoid-related genes in roots but not in shoot tissues. Specifically, SlG1 appears to act together with PSY3 exclusively in the roots to eventually produce SLs but not other carotenoid-derived hormones such as ABA. SLs released by plant roots to the soil are well-known signaling molecules for colonization by AM fungi but also for germination of parasitic plants (Bouwmeester et al., 2021; López-Ráez et al., 2009; Yoneyama et al., 2008). The particular characteristics of the *slg1* mutant make it an attractive target for breeding to negate the negative effects associated to infection by parasitic plants without altering normal shoot growth and metabolism, including photosynthesis and fruit ripening.

## SUPPORTING METHODS

### Methods S1. Metabolite analyses

#### Isoprenoid extraction and quantification

Carotenoids, chlorophylls and tocopherols were extracted as described (Barja et al., 2021; Ezquerro et al., 2022) with some modifications. Freeze-dried tissue powder from roots (25 mg) or leaves (8 mg) was mixed in 2 ml Eppendorf tubes with 375 μl of methanol as extraction solvent, 25 μl of a 10 % (w/v) solution of canthaxanthin (Sigma) in chloroform as internal control, and three 2 mm glass beads. In the case of pericarp tissue, freeze-dried powder (20 mg) was 1 ml of 2:1:1 hexane:acetone:methanol was used instead of methanol. Following extraction as described (Barja et al., 2021; Ezquerro et al., 2022), dry residues were resuspended in 200 μl of acetone by using an ultrasound bath and filtered with 0.2 μm filters into amber-colored 2 ml glass vials. Separation and quantification of individual carotenoids, chlorophylls and tocopherols was performed as described (Barja et al., 2021).

#### ABA extraction and quantification

For ABA quantification, 20 mg of freeze-dried root powder was mixed using 1 ml of 10% methanol in water (containing internal ABA standard) by shaking in a TissueLyser II (Quiagen) at 27 Hz for 3 min. Next, samples were placed in a rotator for 1 hour at 4°C. Extracts were then centrifuged at 14000 x*g* at 4°C for 15 min. Around 800 µl of the liquid phase was recovered in 1.5 ml Eppendorf for further steps. Eluted liquid was run through an Oasis HLB (reverse phase) column as described (Flokova et al., 2014). The dry residues were dissolved in 1 ml of pure methanol and after, dried under a nitrogen flow in a fume hood for 40 min. The eluate was dissolved in 5% acetonitrile and ABA was detected using a reverse phase UHPLC chromatography. The gradient used contains 5 to 50% acetronitrile with 0.05% acetic acid, at a flow speed of 400 μL/min over 21 min. Quantification of ABA was perform with a Q-Exactive mass spectrometer (Orbitrap detector; ThermoFisher Scientific) in conjunction with internal standards (deuterium-labelled hormone at 1pmol/µl), calibration curves and the TraceFinder 4.1 SP1 software.

#### SL extraction and quantification

Tomato seedlings growing in a mixture of river sand and a clay granulate (1:1) were supplied with half-strength Hoagland solution twice a week for 28 days. After 28 days, plants were divided into two groups of which one continued to get the half-strength Hoagland solution for 7 days, while the other was exposed to phosphorus deficiency (by using half-strength Hoagland solutions without phosphate) to induce SL biosynthesis and exudation. Pots were washed with 100 ml of sterile mili-Q water to collect the SLs, which were then concentrated using C18 SPE columns (SUPELCO, Discovery® DSC-18 SPE, 500 mg/3 mL) and analyzed by LC-MS/MS as described (Zhang et al., 2014; 2018).

#### Analysis of volatile organic compounds

For the analysis of VOCs, frozen tomato leaf powder (150 mg) was mixed in a 15 ml glass vial with 1 mL of a saturated CaCl_2_ solution and 100 μL of 750 mM EDTA (pH 7.5). The vial was sealed and sonicated for 5 min and volatile compounds extraction was performed by Head Space Solid-Phase Microextraction (HS-SPME) as previously reported (López-Gresa et al., 2017). VOCs were analyzed in an Agilent 6890N (Santa Clara, CA, USA) gas chromatograph coupled to an Agilent 5975B Inert XL electronic impact (EI) mass detector with an ionization energy of 70 eV and a source temperature of 230°C. Chromatograms were processed using the Enhanced ChemStation software (Agilent). Final identification of VOCs was performed using commercial standards as reported (López-Gresa et al., 2018).

### Methods S2. Gene expression analyses

#### Gene co-expression network (GCN) analyses

The data set used for Gene co-expression network construction was previously described (Wang et al., 2021) and it is publicly available in https://dataview.ncbi.nlm.nih.gov/object/PRJNA679261?reviewer=vs5lk0a94j04c2rgieta1lrlro. It is composed by tomato root samples grown in +P and -P conditions. Tomato isoprenoid biosynthetic genes were retrieved from Plaza 4.0 using Arabidopsis homologs as queries (https://bioinformatics.psb.ugent.be/plaza/) as previously described (Wang et al., 2022). Pairwise pearson correlation coefficients (PCC) were calculated between every two genes using *SlG1*, *SlG2* and *SlG3* as baits and tomato isoprenoid biosynthetic genes as preys. Figures were constructed using R software (https://www.r-project.org/).

#### RNA extraction and RT-qPCR analyses

Total RNA from freeze-dried leaves and roots was extracted using TriPure isolation reagent (Sigma) combined with a Qiagen RNeasy mini spin column kit following manufacturer’s instructions. RNA was quantified using a NanoDropTM 8000 spectrophotometer (ThermoFisher Scientific). ThermoFisher First Strand cDNA synthesis kit using oligo(dT) primer was used to reverse transcribe 1000 ng of RNA into 20 μL of cDNA, which was subsequently diluted 10-fold with mili-Q water and stored at -20 °C for further analysis. Relative mRNA abundance was evaluated via Real-Time Quantitative Polymerase Chain Reaction (RT-qPCR) in a reaction volume of 10 μL containing 5 μL of SYBR Green Master Mix (Thermo Fisher Scientific), 0.3 μM of each specific forward and reverse primer (Table S1) and 2 μL of cDNA. Transcript abundance was evaluated via real-time quantitative PCR (RT-qPCR) in a reaction volume of 10 μl containing 2 μl of the cDNA dilution, 5 μl of SYBR Green Master Mix (Thermo Fisher Scientific), and 0.3 μM of each specific forward and reverse primer (Table S1). The RT-qPCR was carried out on a QuantStudio 3 Real-Time PCR System (Thermo Fisher Scientific) using three independent biological samples and three technical replicates of each sample. Normalized transcript abundance was calculated as previously described (Simon, 2003) using tomato *ACT4* (Solyc04g011500) as endogenous reference gene.

### Methods S3. Co-immunoprecipitation assays

Co-immunoprecipitation experiments were carried out as described (Barja & Rodríguez-Concepción, 2020; Barja et al., 2021) (Methods S3). Constructs encoding Myc- and HA-tagged tomato GGPPS and PSY proteins (Table S2) (Barja et al., 2021) were transformed into *A. tumefaciens* GV3101 strains. A plasmid containing the Arabidopsis phosphoribulokinase protein with a Myc tag (pGWB417_PRK-Myc) was used as negative control. Agroinfiltration of *N. benthamiana* leaves, sample collection, protein extraction and immunoprecipitation of proteins was performed as described (Barja et al., 2021). Myc- and HA-tagged proteins in input and Co-IP samples were detected by immunoblot analyses using 1:2000 αMyc (Sigma) and 1:1000 αHA (Roche) dilutions. Horseradish peroxidase (HRP)-conjugated secondary antibodies against mouse and rat IgGs, respectively were used in a 1:10000 dilution. Amersham ECL Prime Western Blotting Detection Kit (GE Healthcare) was used for detection and the signal was visualized using the Amersham ImageQuant 800 Western blot imaging system.

## Acknowledgments

We thank Mª Rosa Rodriguez-Goberna for technical help with HPLC analyses, Kristýna Floková for the support with ABA measurements, Ernesto Llamas for providing the pGWB417_AtPRK construct, Rodrigo Therezan and Lazaro E.P. Peres (USP, Brazil) for the tomato *ccd7* line, and Albert Ferrer and Laura Gutiérrez for the pDE-Cas9 (with kanamycin resistance) plasmid. This work was funded by grants from Spanish MCIN/AEI/10.13039/501100011033 and European NextGeneration EU/PRTR and PRIMA programs to MR-C (PID2020-115810GB-I00 and UToPIQ-PCI2021-121941). MR-C is also supported by CSIC (202040E299) and Generalitat Valenciana (PROMETEU/2021/056). ME received a predoctoral fellowship from MCIN/AEI (BES-2017-080652) and an EMBO Scientific Exchange Grant (9315). No conflict of interest is declared.

## Author Contributions

ME and MR-C designed the research; ME, CL, MVB, EB-E, JP-P and YW performed research; LD, MPL-P, PL and HB contributed new analytic tools; ME, CL, MVB, EB-E, JP-P, YW, LD, PL, MPL-G, HB and MR-C analyzed and discussed data; ME and MR-C wrote the paper and incorporated the input of the rest of the authors.

## Data availability

The data that supports the findings of this study are available in the supplementary material of this article.

---

**Figure S1.**
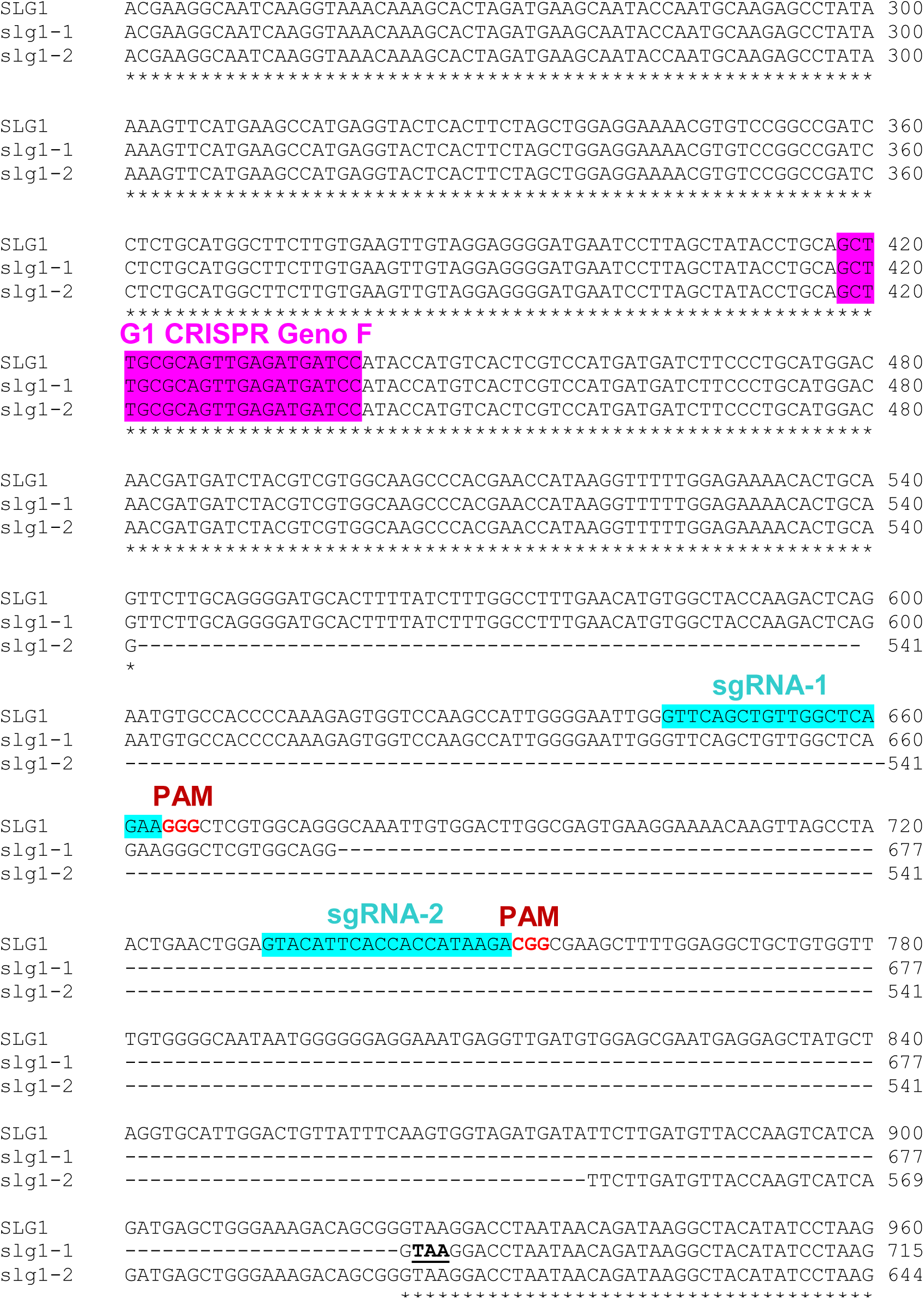
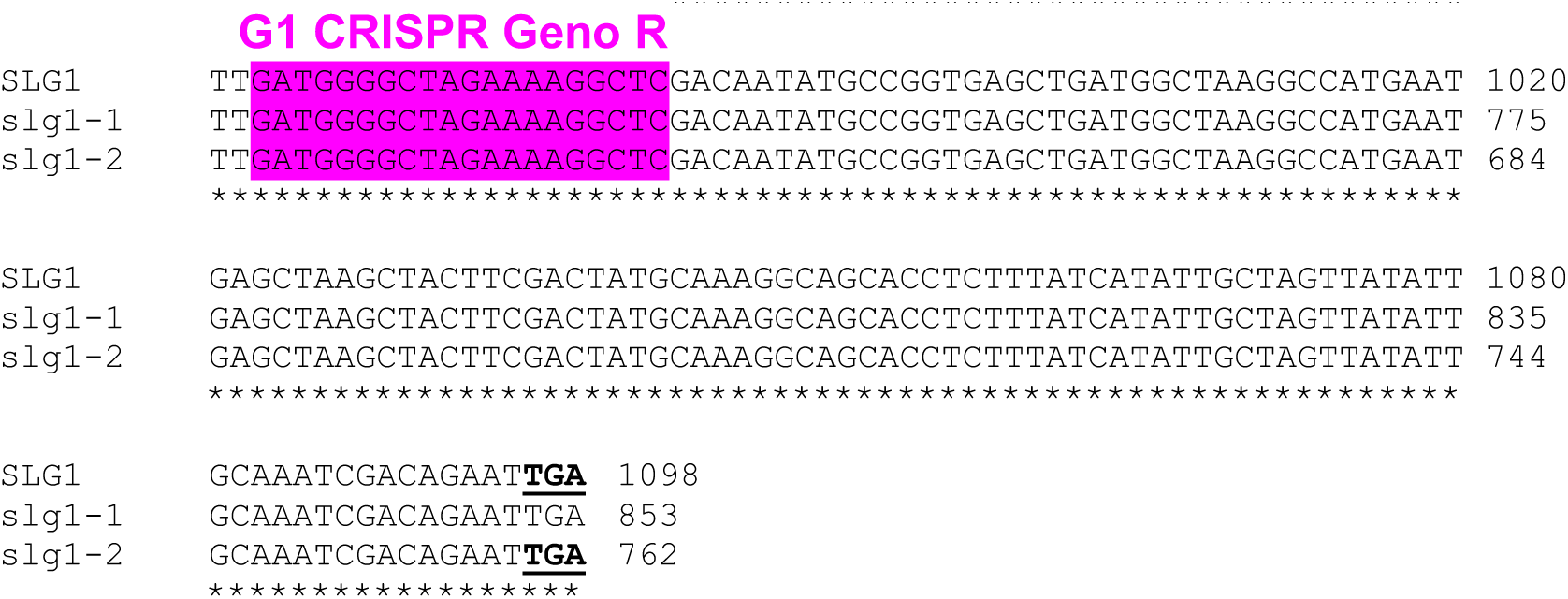
DNA sequence alignment of *SlG1* CRISPR mutants. Alignment was performed using Clustal Omega (https://www.ebi.ac.uk/Tools/msa/clustalo/) with default settings. The sequence encoding the predicted plastid-targeting peptide is marked in green. Designed single-guide RNAs (sgRNA) are highlighted in blue and genotyping oligonucleotides are highlighted in fucsia. Protospacer adjacent motifs (PAM) are highlighted in red. Translation stop codons are underlined and marked in bold. Numbers at the end of each sequence indicate DNA sequence length.

**Figure S2.**
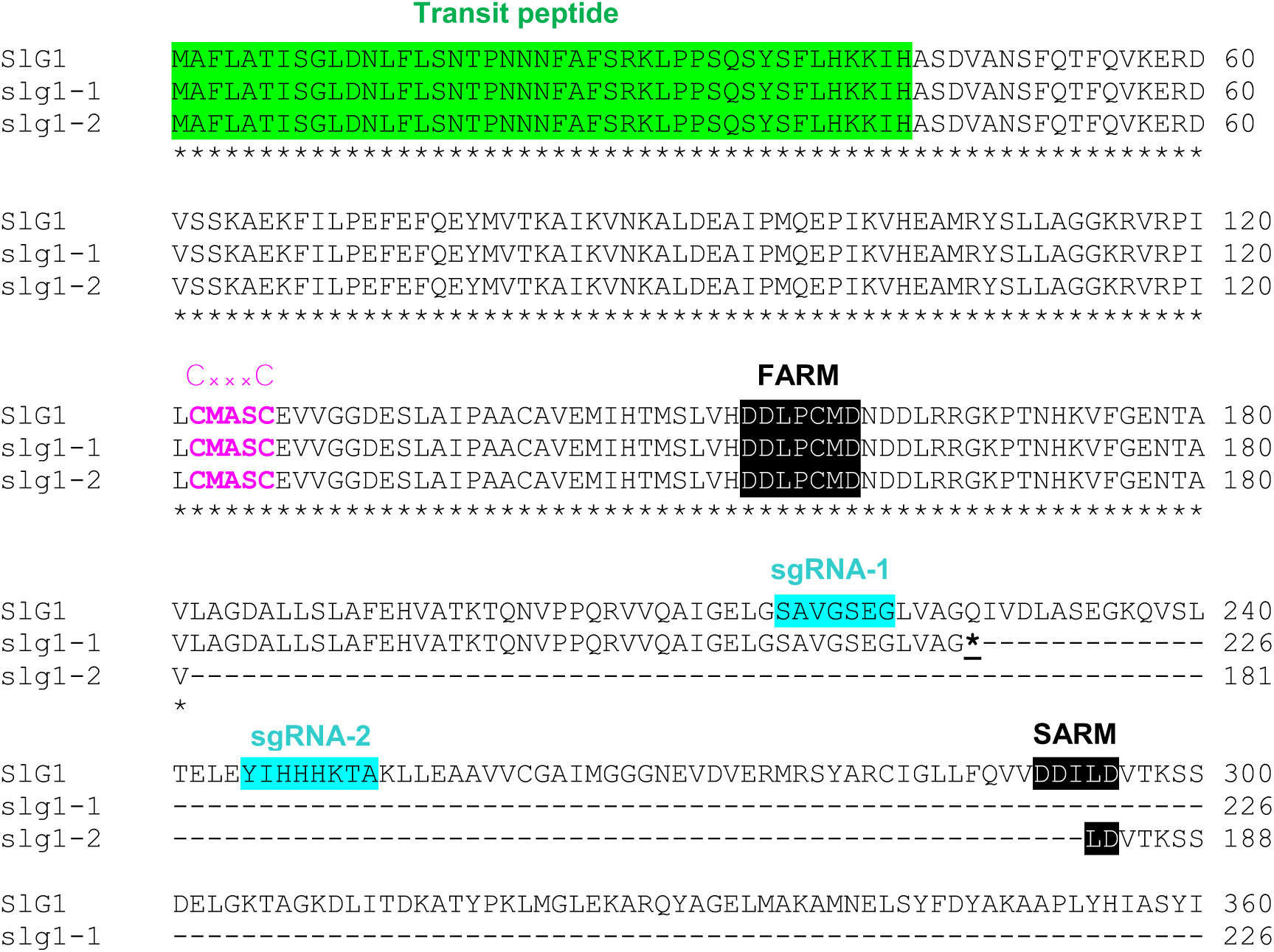
Protein alignments of SlG1 wild-type sequences with the selected CRISPR mutants. Multiple sequence alignment was performed using *Clustal Omega* (https://www.ebi.ac.uk/Tools/msa/clustalo/) with default settings. The predicted targeting peptides, the region of the designed sgRNAs and the catalytic motifs FARM (first aspartate-rich motif) and SARM (second-aspartate rich motif) are boxed in green, blue and black, respectively. The protein-protein interaction CxxxC (x = any hydrophobic residue) motifs are highlighted in pink. Numbers at the end of each sequence indicate protein length.

**Figure S3.**
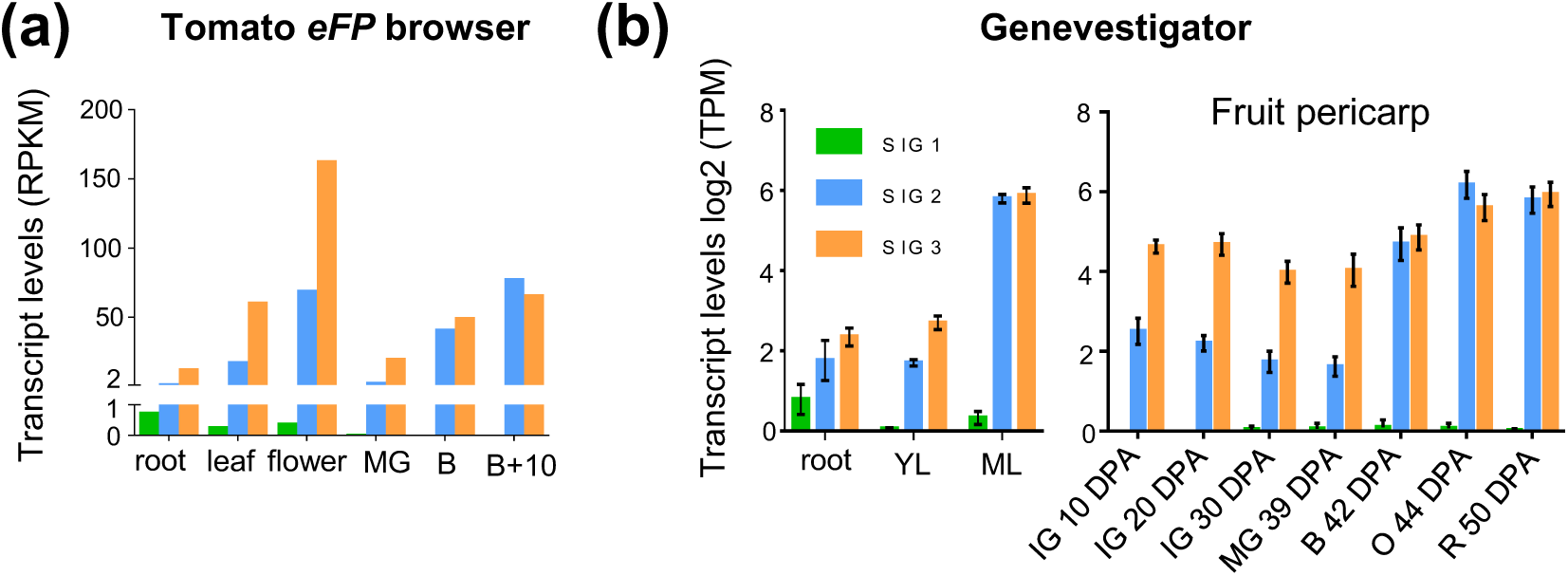
*SlG1, SlG2 and SlG3* transcript levels in different tissues and developmental stages. **(a)** RNAseq data retrieved from the *Tomato eFP Browser* database (http://bar.utoronto.ca/efp_tomato/cgi-bin/efpWeb.cgi). Levels are represented as RPKM (Reads per Kilobase of transcript per Million mapped reads). **(b)** RNAseq data obtained from Genevestigator (https://genevestigator.com). Levels are represented as log2 TPM (Transcripts Per Million mapped reads). Abbreviations: DPA, days post-anthesis; IG, immature green; MG, mature green; B, breaker; O, orange, R, red. YL, young leaves; ML, mature leaves.

**Figure S4.**
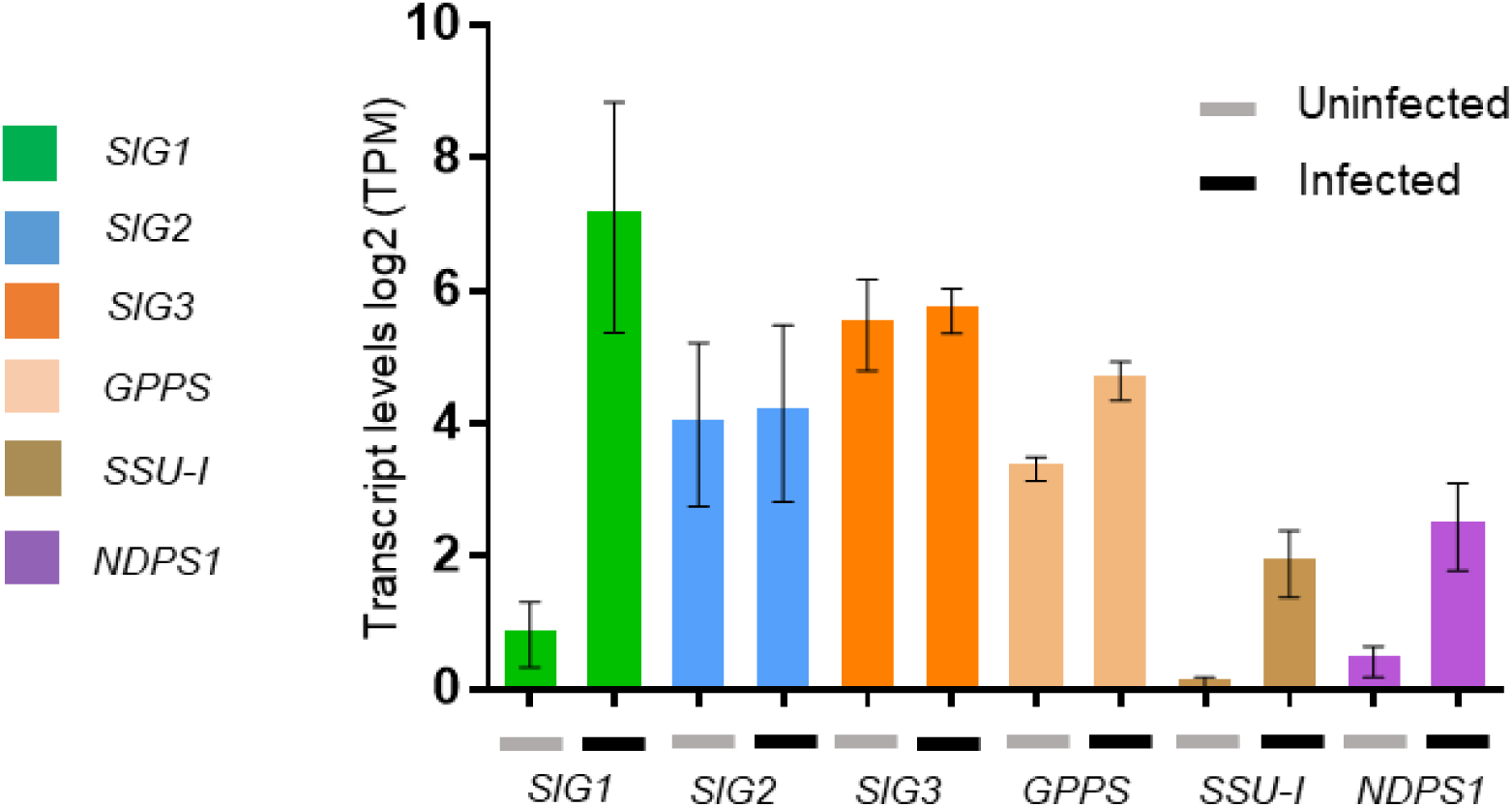
Expression levels of *SlGGPPS, SlGPPS, SlSSUI and SlNPS1* in leaves after *Pseudomonas syringae* infection. RNAseq data obtained from Genevestigator (https://genevestigator.com) of three different tomato infection experiments with *P.syringae pv. tomato* DC3000. Two different tomato cultivars were used, Ailsa Craig (Yang et al. 2016) and Rio Grande (Rosli et al., 2013; Pombo et al., 2014). Samples were collected at 9 (Ailsa Craig) or 6 (Rio Grande) hours post infection, respectively. Plots show the transcript levels of the indicated genes in leaves of both cultivars during *P. syringae pv. tomato* DC3000 infection and are shown as log2 TPM (Transcripts per Million mapped reads).

**Figure S5.**
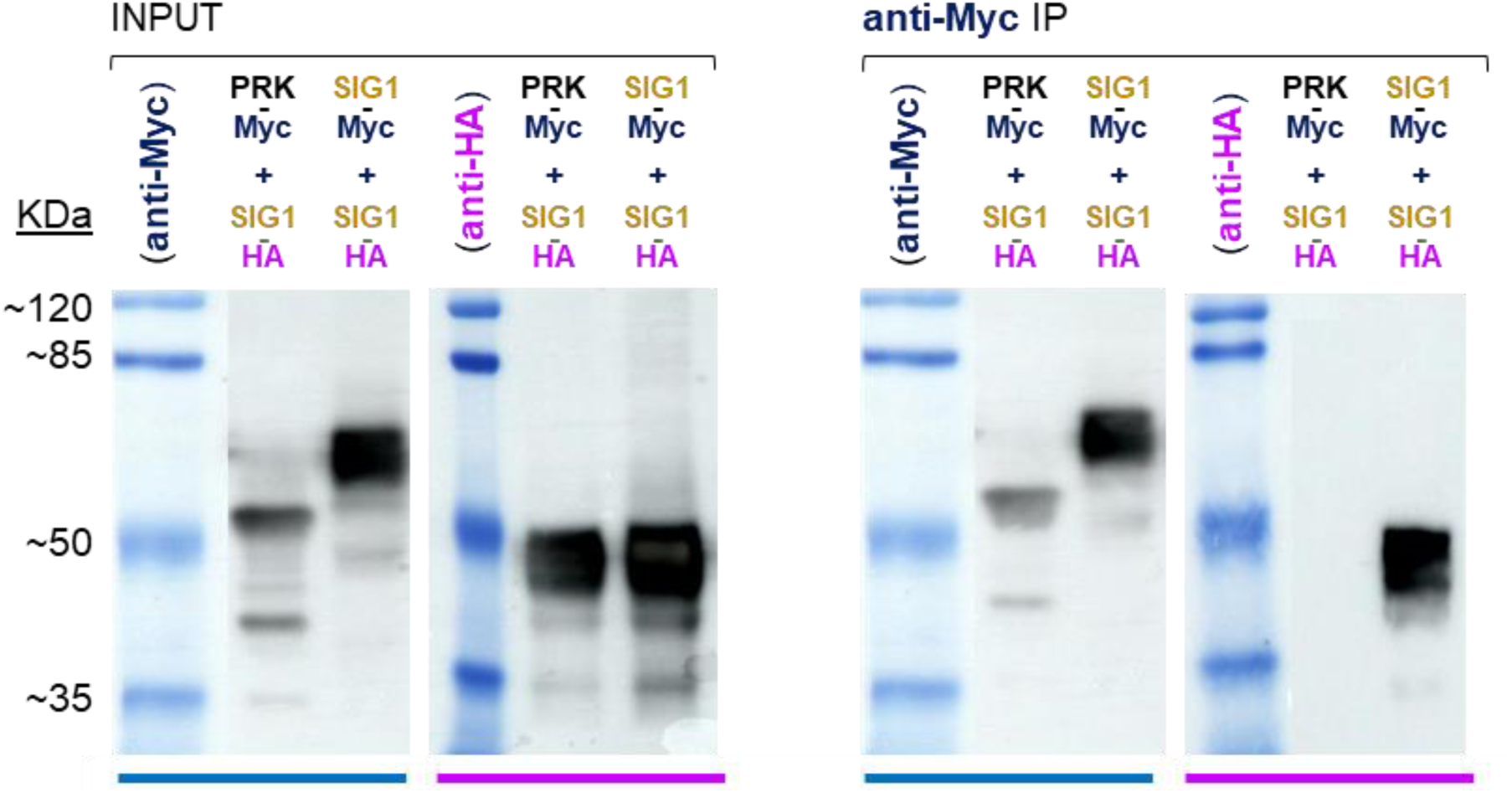
SlG1-Myc is able to specifically bind to protein partners. *N. benthamiana* leaves were co-agroinfiltrated with constructs encoding the indicated proteins tagged with C-terminal Myc or HA epitopes. Immunoblot analysis of crude extracts (INPUT) with anti-Myc (dark blue) and anti-HA (purple) antibodies was carried out to confirm successful protein production. The same samples were used for immunoprecipitation (IP) with anti-Myc followed by immunodetection with anti-Myc (to confirm successful IP) or anti-HA (to identify co-immunoprecipitated partners). Predicted protein molecular weights (KDa): PRK-Myc, 52.1; SlG1-Myc, 56.9; SlG1-HA 47.8.

**Table S1.**
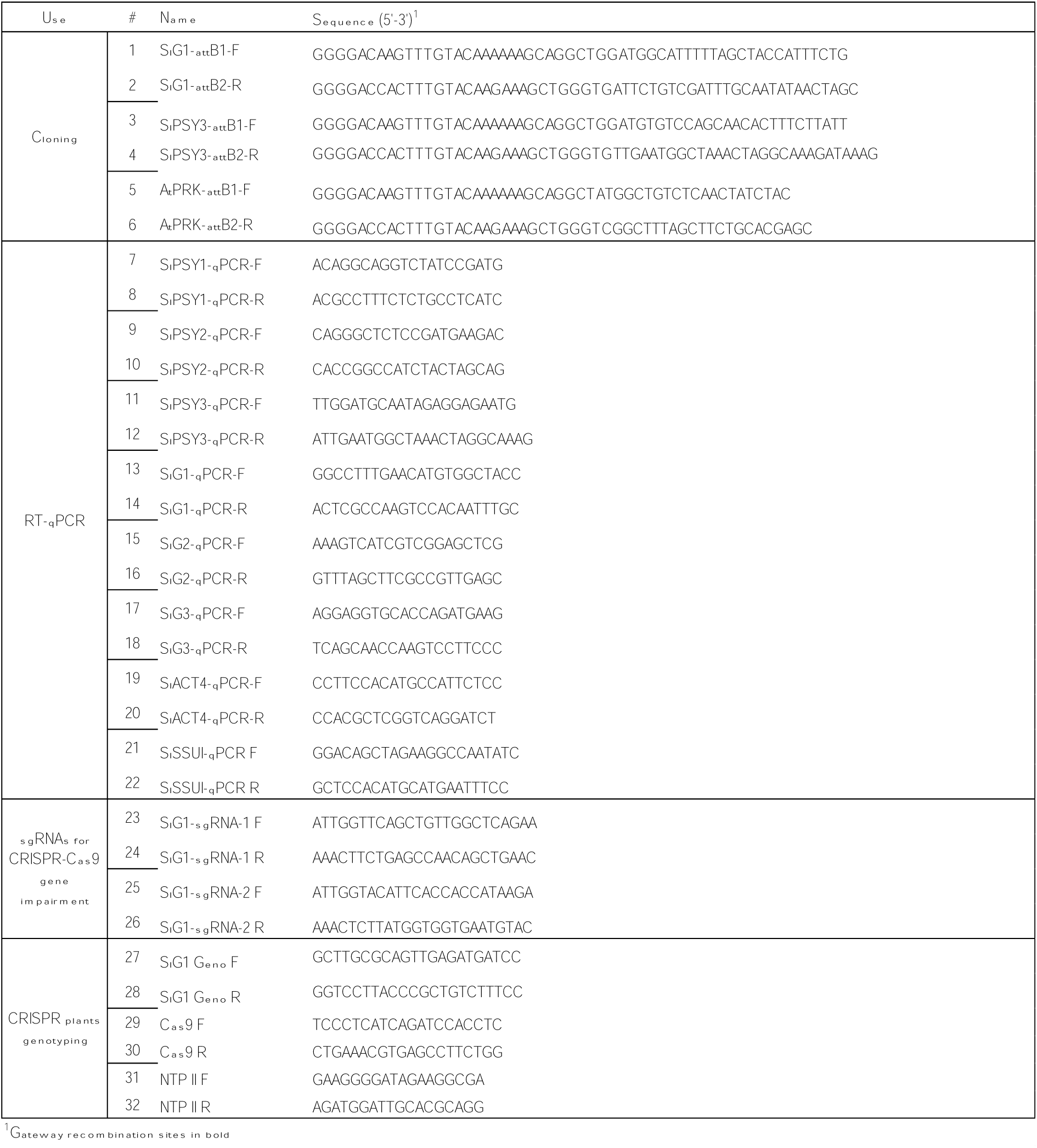
List of primers used in this work

**Table S2.**
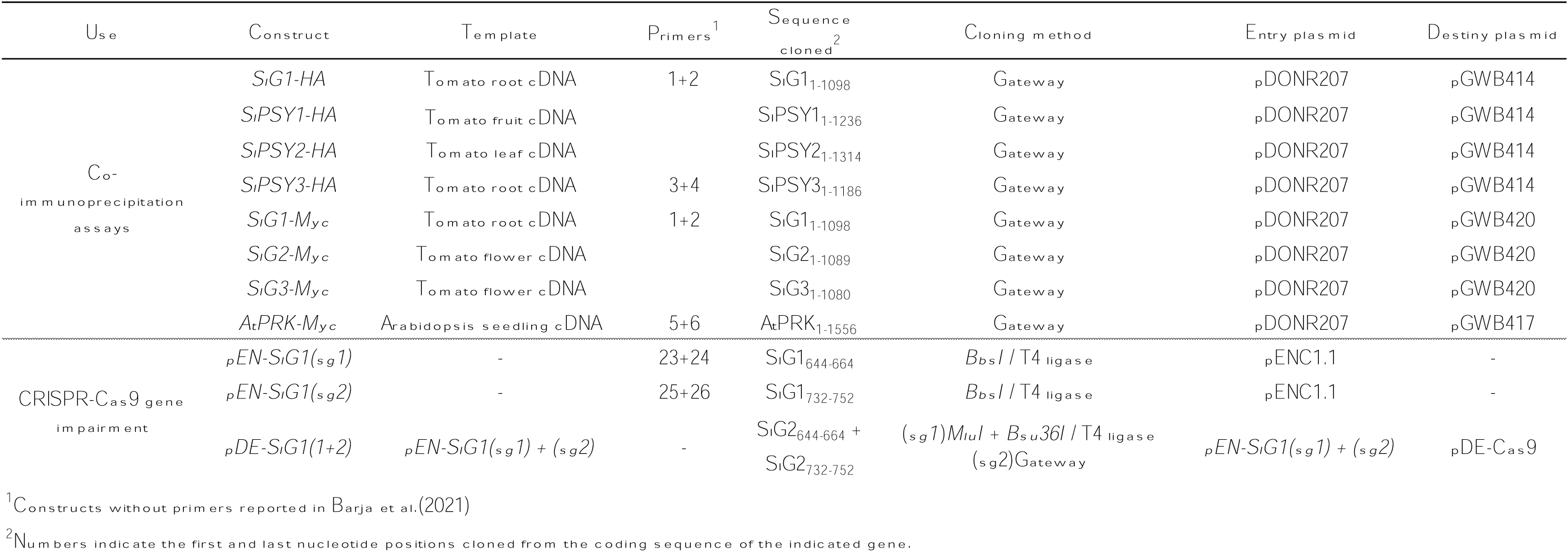
Constructs and cloning details.

**Table S3.**
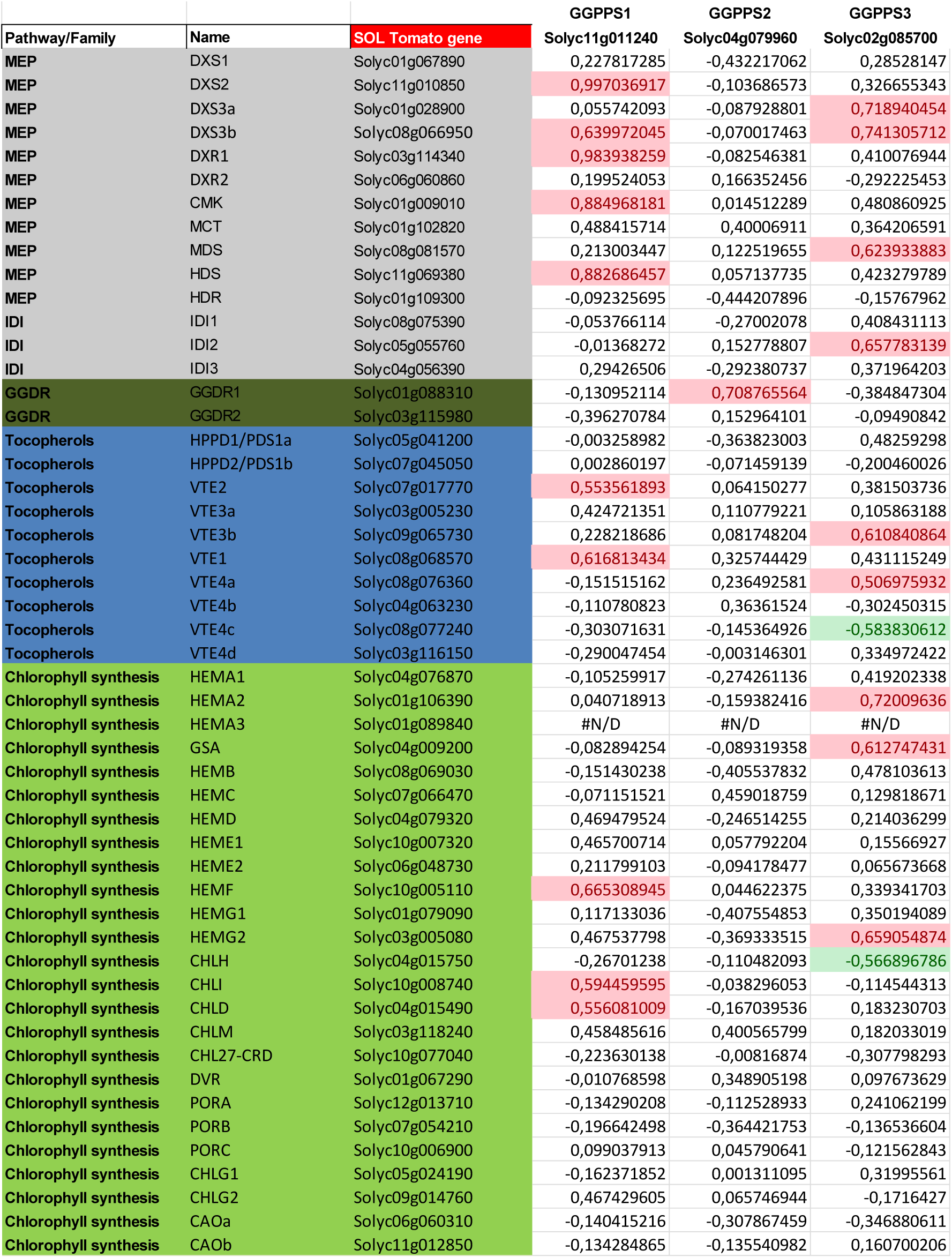
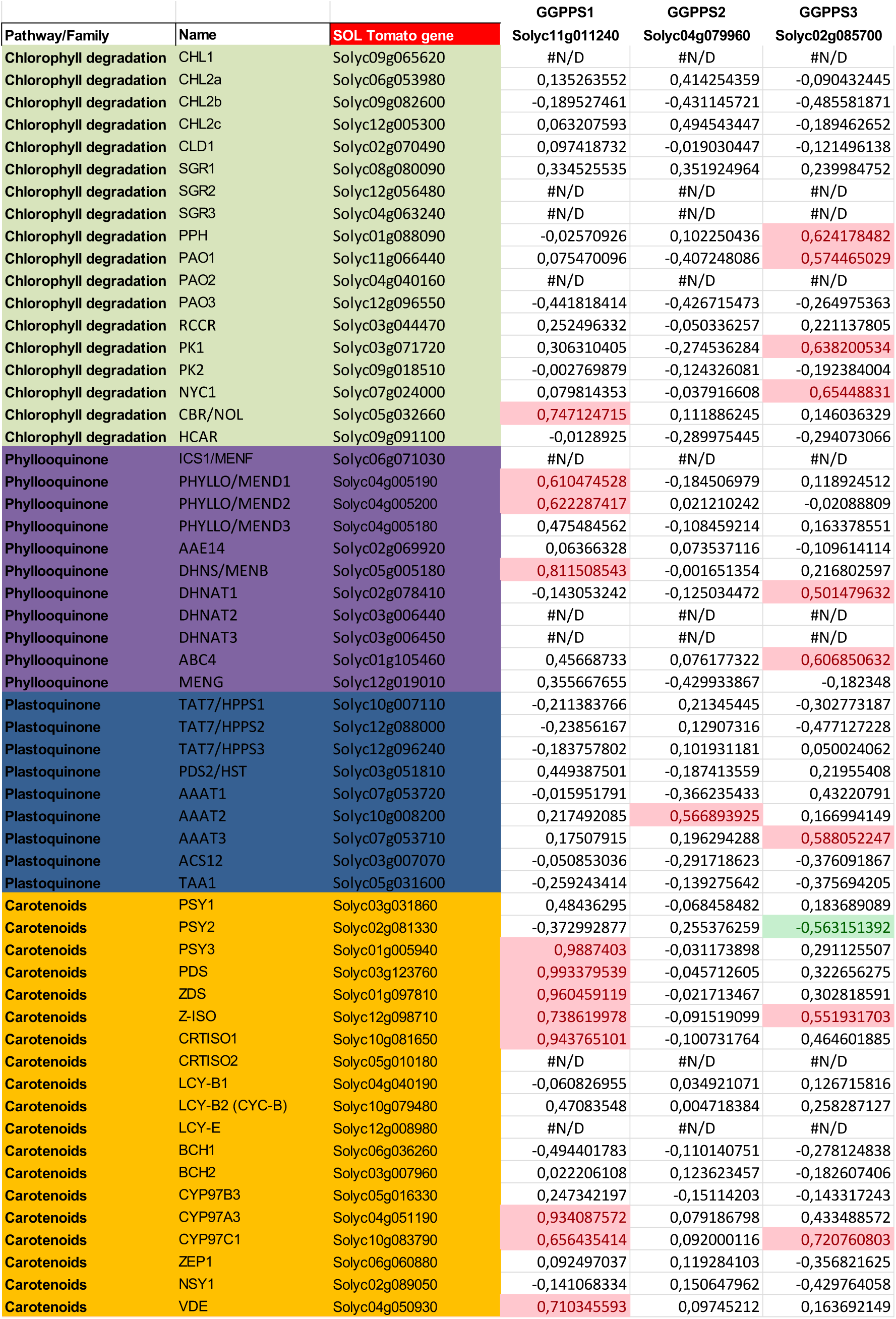
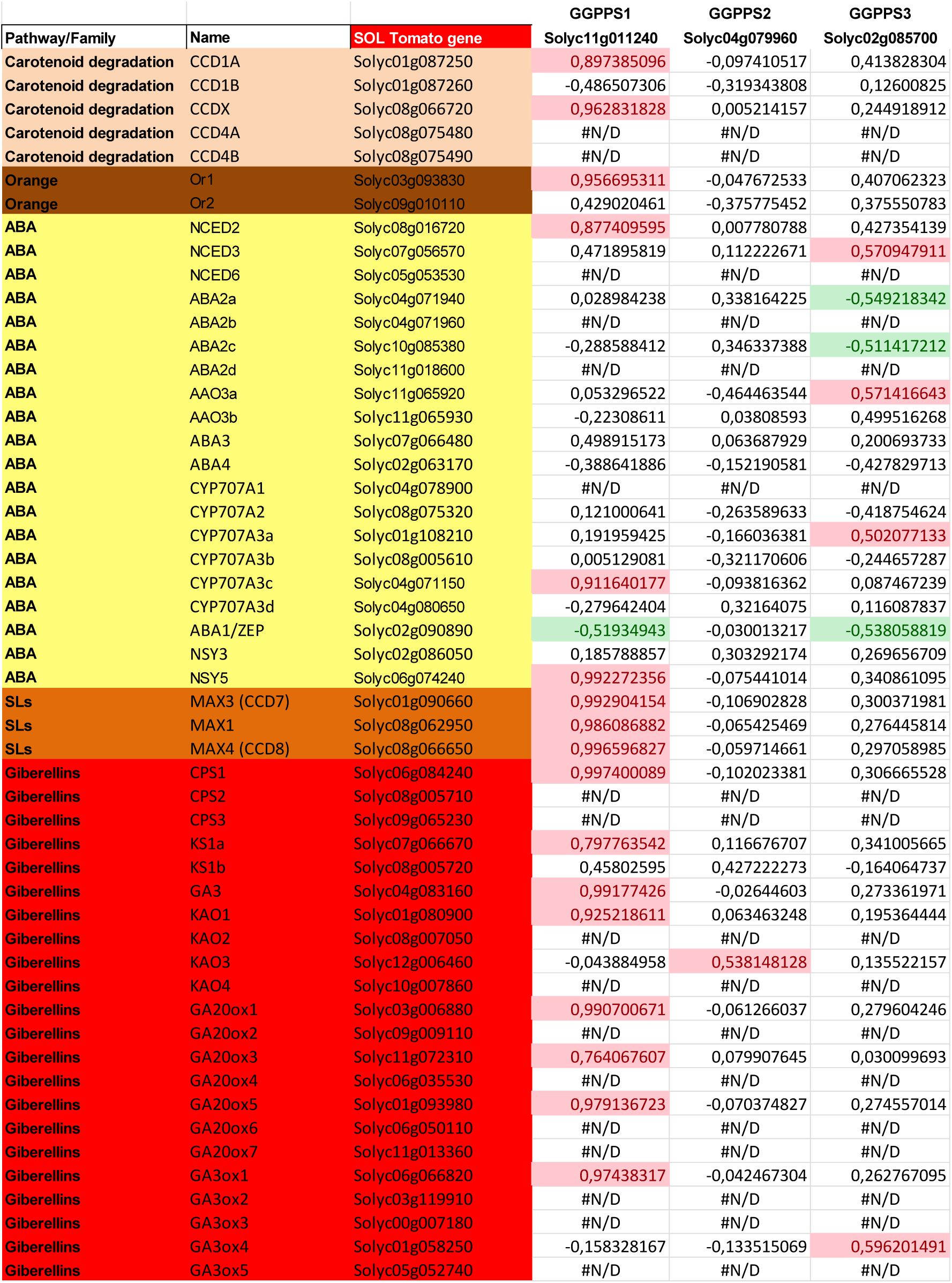
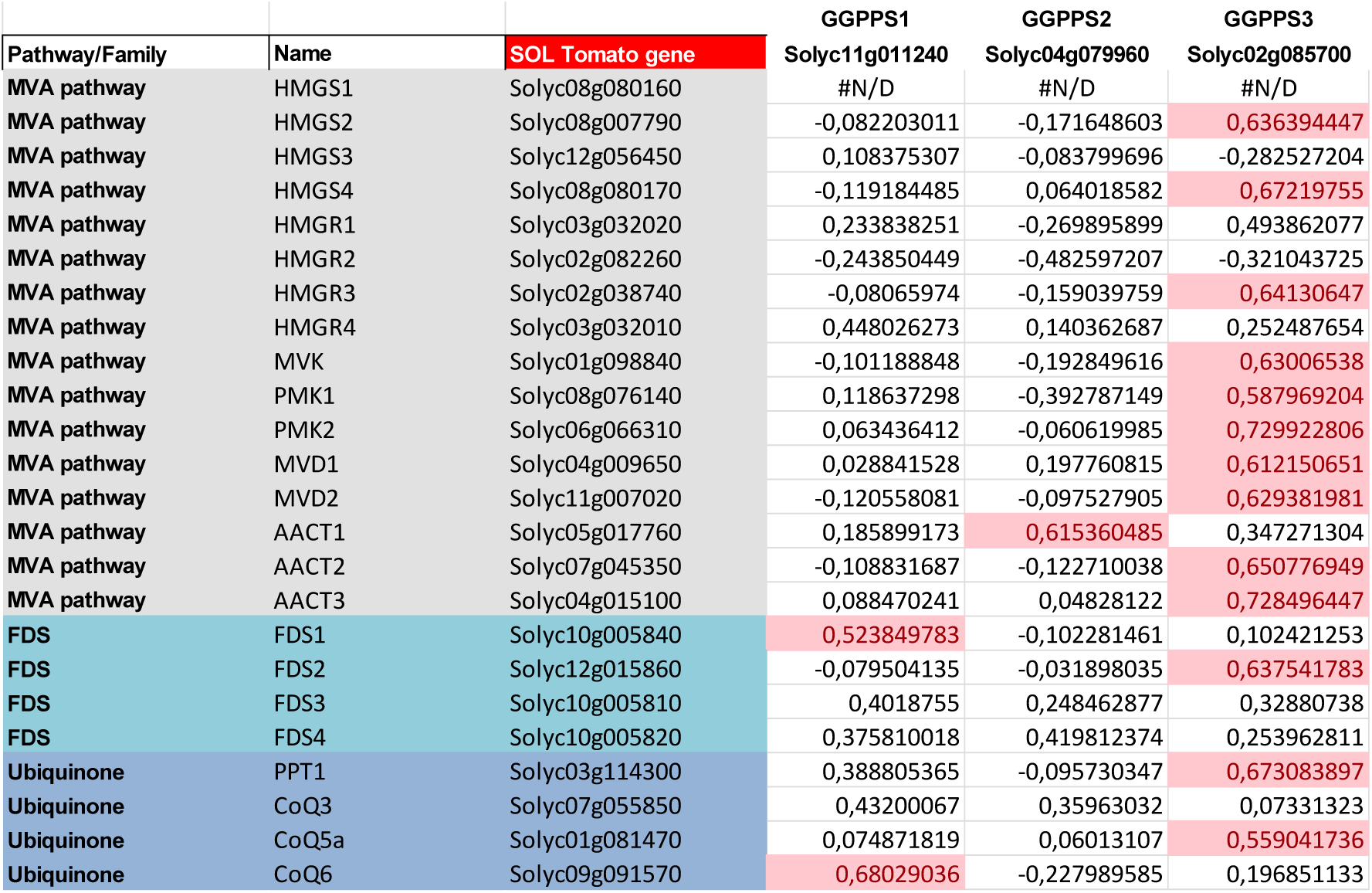
Co-expression of tomato GGPPS paralogs (guide genes) with isoprenoid-related genes (query genes) in root tissue. Significant pairwise Pearson correlations between guide and query genes (≥0.55) are highlighted in red (when positive) or green (when negative). Arabidopsis genes were used as queries to search for tomato homologs in PLAZA 4 (https://bioinformatics.psb.ugent.be/plaza/versions/plaza_v4_dicots/). Genes are organized by pathways.

